# Microbial education plays a crucial role in harnessing the beneficial properties of microbiota for infectious disease protection in *Crassostrea gigas*

**DOI:** 10.1101/2024.05.17.594303

**Authors:** Luc Dantan, Prunelle Carcassonne, Lionel Dégremont, Benjamin Morga, Marie-Agnès Travers, Bruno Petton, Mickael Mege, Elise Maurouard, Jean-François Allienne, Gaëlle Courtay, Océane Romatif, Juliette Pouzadoux, Raphaël Lami, Laurent Intertaglia, Yannick Gueguen, Jeremie Vidal-Dupiol, Eve Toulza, Céline Cosseau

**Affiliations:** IHPE, Univ. Montpellier, CNRS, IFREMER, Univ. Perpignan Via Domitia, Perpignan, France; Ifremer, ASIM, F-17390 La Tremblade, France; IHPE, Univ. Montpellier, CNRS, IFREMER, Univ. Perpignan Via Domitia, Montpellier, France; Univ Brest, Ifremer, CNRS, IRD, LEMAR, F-29280 Plouzané, France; Sorbonne Université, CNRS, Laboratoire de Biodiversité et Biotechnologies Microbiennes, Observatoire Océanologique de Banyuls-sur-Mer, Avenue Pierre Fabre, 66650, Banyuls-sur-Mer, France; Sorbonne Université, CNRS, Fédération de Recherche, Observatoire Océanologique, 66650 Banyuls-sur-mer, France; MARBEC, Univ Montpellier, CNRS, Ifremer, IRD, Sète, France

**Keywords:** *Crassostrea gigas*, Microbial education, Oyster holobiont, OsHV-1 µVar, *Vibrio aestuarianus*

## Abstract

**Background:** Recently, the frequency and severity of marine diseases have increased in association with global changes, and molluscs of economic interest are particularly concerned. Among them, the Pacific oyster (*Crassostrea gigas*) production faces challenges from several diseases such as the Pacific Oyster Mortality Syndrome (POMS) or vibriosis. Various strategies such as genetic selection or immune priming have been developed to fight some of these infectious diseases. The microbial education, which consist of exposing the host immune system to beneficial microorganisms during early life stages is a promising approach against diseases. This study explores the concept of microbial education using controlled and pathogen-free bacterial communities and assesses its protective effects against POMS and *Vibrio aestuarianus* infections, highlighting potential applications in oyster production.

**Results:** We demonstrate that it is possible to educate the oyster immune system by adding microorganisms during the larval stage. Adding culture based bacterial mixes to larvae protects only against the POMS disease while adding whole microbial communities from oyster donors protects against both POMS and vibriosis. The efficiency of the immune protection depends both on oyster origin and on the composition of the bacterial mixes used for exposure. No preferential protection was observed when the oysters were stimulated with their sympatric strains. We further show that the added bacteria were not maintained in the oyster microbiota after the exposure, but this bacterial addition induced long term changes in the microbiota composition and oyster immune gene expression.

**Conclusion:** Our study reveals successful immune system education of oysters by introducing beneficial micro-organisms during the larval stage. We improved the long-term resistance of oysters against critical diseases (POMS disease and *Vibrio aestuarianus* infections) highlighting the potential of microbial education in aquaculture.

## Introduction

The Pacific oyster *Crassostrea gigas* (also known as *Magallana gigas*) stands as the most widely cultivated oyster species in the world, underpinning a substantial proportion of the aquaculture industry (Food and Agriculture Organisation 2022). However, the production of *C. gigas* faces significant challenges due to recurring infectious diseases, inflicting high mortalities each year (Friedman et al. 2005; Cotter et al. 2010; Pernet et al. 2012; Azéma et al. 2015). Two prevalent infections – the Pacific Oyster Mortality Syndrome (POMS) caused by the Ostreid herpesvirus type 1 µVariant (OsHV-1 µVar) and vibriosis initiated by *Vibrio aestuarianus* infection *−* are primarily responsible for these alarming mortalities. POMS is a complex and polymicrobial disease which preferentially affects younger oysters and can decimate up to 100% of the spat in French farms (Segarra et al. 2010; Petton et al. 2021). The infection by OsHV-1 µVar marks a critical step in the progression of POMS, inducing an immunocompromised state in oysters by altering haemocytes physiology (de Lorgeril et al. 2018; Petton et al. 2021). This leads to a dysbiosis of oyster microbiota and results in colonisation by opportunistic bacteria and death of the oyster (de Lorgeril et al. 2018; King et al. 2019a; Petton et al. 2021). On the other hand, *V. aestuarianus* is another harmful primary pathogen with chronic mortality reaching a cumulative mortality rate up to 30%. This loss induces important economic consequences since it preferentially infects market size oysters which have been raised for several years (Azéma et al. 2017; Lupo et al. 2019).

Efforts to combat these infectious diseases have spawned various approaches based on the increasing knowledge and resources available on oysters. Genetic selection is a promising avenue which aims at selecting pathogen-resistant oysters (Dégremont et al. 2015, 2020). However, this approach exhibits limitations such as the potential selection of trade-offs which could counter select traits important for the commercial value of *C. gigas*. Moreover, the demonstration of the existence of immune priming in *C. gigas* has opened up a whole new field of applications based on the use of viral mimics (Lafont et al. 2017, 2020; de Kantzow et al. 2023; Montagnani et al. 2024). However, this innovative approach only protects against POMS infections (Green and Montagnani 2013; Lafont et al. 2017). A diversity of studies on oyster-microbiota interactions have also opened a new field of investigations consisting in identifying bacteria beneficial for their associated host during adverse conditions (King et al. 2019a; Clerissi et al. 2020; Delisle et al. 2022; Fallet et al. 2022). Research on disease prevention in molluscs based on the use of probiotics has been ongoing for decades but has yet to see widespread applications in farms (Yeh et al. 2020; Takyi et al. 2023, 2024; Muñoz-Cerro et al. 2024). While several pre/probiotic-based methods to mitigate infectious diseases have demonstrated success in shrimp hatcheries (Swain et al. 2009; Pham et al. 2014; Wen et al. 2014), their application in oyster farming, particularly in open-sea environments, faces distinct challenges and limitations. Oysters, as filter-feeding organisms, often face complex microbial interactions in their natural habitats (Lokmer et al. 2016). Consequently, achieving and maintaining a precise balance of beneficial microorganisms through probiotics addition can be challenging. Additionally, their culture in open-sea present limitations in the implementation of probiotics.

The concept of microbial education, consists in exposing the host immune system to beneficial microorganisms during early development (Arrieta et al. 2014; Gensollen et al. 2016). This is because early life stages represent critical periods of growth and development where the host’s immune system is still maturing (Renz et al. 2017). This strategy offers significant advantages on oysters, as it can confer a protective effect while allowing exposure in hatchery during the larval phase in controlled environments (Dantan et al. 2024). Numerous studies have shown that a proper host-microbiota interaction during the early development plays an important role in the long term host immune responses in a wide range of marine organisms (Chung et al. 2012; Galindo-Villegas et al. 2012; Abt and Artis 2013; Sommer and Bäckhed 2013). In this context, Fallet and colleagues (Fallet et al. 2022) explored the potential of using wild-microbiota to educate the immune system of *C. gigas*. Through a ten-day exposure of *C. gigas* larvae to a whole microbiota from donor oysters, they induced a long-term beneficial effect. The microbiota-exposed oysters exhibited enhanced resistance to OsHV-1 µVar, resulting in improved survival rates compared to non-exposed counterparts. This study underscored the crucial role of microbiota on oyster immune system education, suggesting potential applications in commercial hatcheries. However, concerns regarding exposure to hazardous uncontrolled microbial communities transferred from donor oysters necessitate a cautious approach as it might contain primary or opportunistic pathogens. Indeed, prior to the recipient larvae exposure performed in Fallet *et al*. study, the donor oysters were placed in farming area during a non-infectious period to allow oysters to capture the maximum diversity of field microorganisms. Then, these donor oysters were placed in the rearing tanks during larval development where they transmitted their highly diverse microbial community to the recipient larvae. Although the donor oysters were considered healthy (Le Roux et al. 2016; Fleury et al. 2020), the presence of undetectable pathogens cannot entirely be excluded

Here, our study aimed to explore the feasibility of microbial education in oyster larvae while considering and mitigating the risks associated with uncontrolled transfer of hazardous microorganisms found in wild-microbiota. We investigated whether exposing oyster larvae to a controlled, pathogen-free bacterial community from donor oysters that had always been maintained in biosecured facilities could confer the protective effects against POMS and *V. aestuarianus* infection. Additionally, we examined the feasibility of microbial education using a reduced synthetic bacterial community composed of cultivable bacteria isolated from disease resistant oysters. For this purpose, we developed and tested multi-strain bacterial mixes originating from the same geographical areas as the recipient oyster populations used in this study. Our comprehensive assays encompassed three distinct oyster populations from the Atlantic Ocean (Brest bay, La Tremblade in Marennes-Oleron bay, and Arcachon bay) and one from the Mediterranean Sea (Thau lagoon), enabling an in-depth exploration of the potential differential effects of bacterial exposure to either sympatric or allopatric oyster populations.

## Materials and methods

### Oyster sampling

Oysters were collected along the French Atlantic coasts, during two different sampling campaigns (in February 2020 and November 2020), while it was only in November 2020 for the Mediterranean site due to covid restrictions arisen earlier in the year. For the Atlantic coast, 3 sites were selected: the Brest bay (Brittany, France; lat 48.3349572; long –4.3189134), La Tremblade in Marennes-Oleron bay (Nouvelle-Aquitaine, France; lat 45.8029675; long – 1.1534223) and the Arcachon bay (Nouvelle-Aquitaine, France; lat 44.6813750; long – 1.1402178). For the Mediterranean coast, the selected site was the Thau lagoon (Occitanie, France; lat 43.39404; long 3.58092). For each site, 5 oysters (average weight = 2.5 g) were randomly sampled. Hence, the sampled oysters were located on sites with a high density of oysters (wild and farmed) and have therefore survived an annual infectious episode of POMS allowing us to assume they were resistant to the disease but also in the window of permissiveness for *Vibrio aestuarianus* infection (Azéma et al. 2016). Based on these facts, we hypothesized that sampling bacteria from these disease-resistant oysters increases the likelihood of isolating beneficial bacteria.

### Isolation of cultivable bacteria from *Crassostrea gigas*

The five disease resistant oysters sampled on each site were carefully brushed and washed to remove the sediments, epiphytes and epibionts present on the shell. The flesh of the animals was then individually crushed with an Ultra-Turrax T25 mixer (5 x 5 sec) in 15 ml falcon tubes. The homogenized tissues were then diluted at 1:10, 1:100 and 1:1000 in sterile artificial seawater. A hundred µL of each dilution were spread on two Marine Agar (MA) (Marine Agar Difco 2216) plates and incubated at 15°C or 20°C.

After a minimum incubation period of 3 days, bacterial colonies were selected according to their morphotypes. A maximum of different morphotypes were selected to maximise the biodiversity in our sampling and isolated by streaking a colony on a new MA plate and purified by two successive subculturing. Then, the pure cultures of individual bacteria were transferred onto Marine Broth (MB) tube (Marine Broth Difco 2216) at 15°C or 20°C and under a constant agitation. After 48h of growth, 500 µL of these cultures was used for cryopreservation in 35% glycerol (V/V) and put into a –80 °C freezer. About 1 ml of the liquid culture was pelleted for further DNA extraction.

### DNA extraction and identification of the cultivable bacteria

DNA extraction of the bacterial strains isolated from oysters and cultivated on agar plates was carried with the Wizard® Genomic DNA Purification Kit (Promega) according to the manufacturer instructions. 16S rRNA gene sequencing was performed on these samples to identify each bacterium from the collection. The PCR and 16S rRNA gene sequencing was performed by the Genoscreen sequencing facilities (http://www.genoscreen.fr/fr/). Briefly, two pairs of primers P8/PC535 (P8 5’-AGAGTTTGATCCTGGCTCAG; PC535 5’-GTATTACCGCGGCTGCTGGCAC) and 338F/1040R (338F 5’-CTCCTACGGGAGGCAG; 1040R 5’-GACACGAGCTGACGACA) were used for the PCR to amplify the V1-V3 and V3-V5 of the 16S rRNA gene. PCR products were then purified with Sephadex-G50 gel (GE Healthcare) before analysis into ABI 3730XL capillary sequencer. The resulting sequences were then assembled by using the DNA baser sequence assembly software (v4) (Heracle BioSoft, www.DnaBaser.com) and then added in the Ezbiocloud database (Yoon et al. 2017) in order to identify the taxonomy of the isolated bacteria composing the collection.

### Larval cytotoxic effect

Two days old larvae (D stage) were distributed in wells of a 6-well plate filled with three ml of sterile seawater at a density of 10 larvae per ml and maintained at a temperature of 20°C and a 12:12 day:night photoperiod. Treatment (bacterial challenge with a single bacterial strain) and control (only sterile seawater) was each conducted in duplicate. The bacteria were cultivated from glycerol stock in 10 ml of Marine Broth (MB) for 24h at 20°C and then, 1 ml of each bacterial culture was inoculated into 10 ml fresh MB media and incubated at 20°C under constant agitation. After 48 hours of incubation, the OD_600_ was measured, and the right amount of bacteria was collected before being centrifuged at 4000 rpm for 2 minutes and the supernatant was discarded. The pellets were then resuspended in 10 ml sterile seawater. Larvae were challenged by addition of a target concentration of 10^7^ CFU/ml of each bacterial strain (Multiplicity of infection = 10^6^ bacteria per larvae). Larval mortality was recorded 48h post addition of bacteria by evaluation of active swimming and/or gut and cilia movement under binocular microscope.

### Multi-strain bacterial mixes preparation for interaction with oysters

Five multi-strain bacterial mixes were tested (**Table 1**): four site-specific multi-strain bacterial mixes composed of bacteria isolated from oysters sampled at each geographical site (Brest mix, La Tremblade mix, Arcachon mix and Thau mix) and a multi-site bacterial composed of bacteria isolated from oysters sampled on all the different sites. The bacteria were cultivated from glycerol stock in 10 ml of Marine Broth (MB) for 24h at 20°C and then, 1 ml of each bacterial culture was inoculated into 50 ml fresh MB media and incubated at 20°C under constant agitation. After 48 hours of incubation, the OD_600_ was measured, and a quantity of 3.10^8^ CFU was collected and pooled into a same mix for each cultivated bacterium. The mixes were then centrifuged at 4000 rpm for 2 minutes and the supernatant was discarded. The pellets were then resuspended in 10 ml sterile seawater and added immediately to 30 L larval rearing tanks to a final concentration of 10^4^ CFU/ml for each bacterium.

**Table 1:**
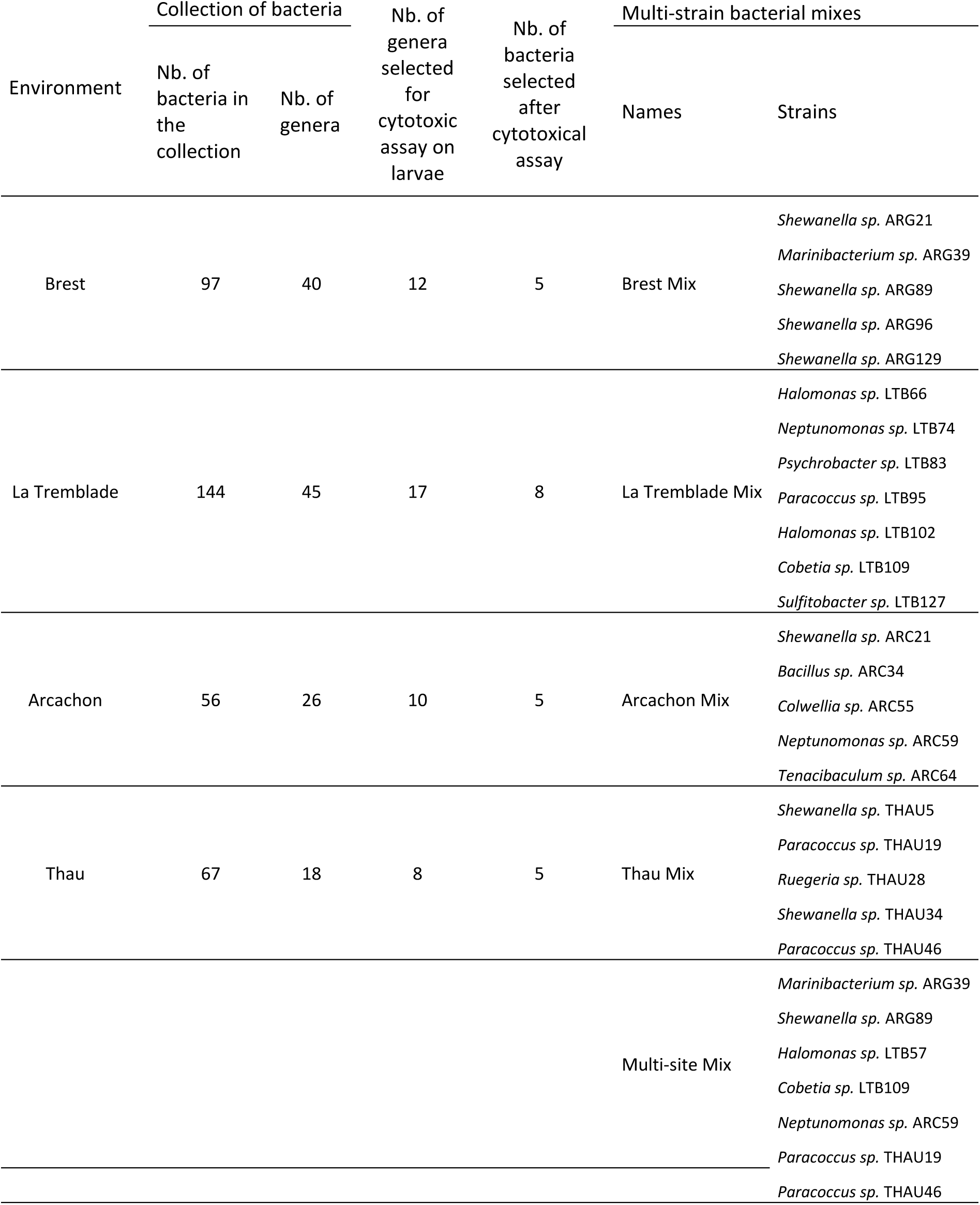
Composition of the 5 multi-strain bacterial mixes produced according to their predictive beneficial properties.

### Oyster reproduction

150 wild oysters were randomly sampled from each geographic site as described above (Brest bay, La Tremblade in Marennes-Oleron bay, Arcachon bay, Thau lagoon) in order to generate 4 oyster populations (Brest, La Tremblade, Arcachon and Thau populations) accordingly to commercial oyster hatchery practices. Briefly, oyster genitors were transferred into the Ifremer hatchery facility in La Tremblade. To avoid eventual horizontal transmission of pathogens among populations, each was placed in separate tanks of 250 L in a flow through system with a water circulation of 500 L/h. Seawater temperature was gradually increased from 10 to 20°C within one week and maintain to 20°C to favour the gametogenesis. Broodstock were fed *ad libitum* with a mixture of phytoplankton (*Isochrysis galbana, Tetraselmis suecica,* and *Skeletonema costatum*). After 2 months, oysters were shucked and sexed by microscopic observation. Only fully mature oysters were used, representing between 20 to 23 genitors per population **(Supplementary File 1, Table S1)**. Spermatozoa and oocytes were collected by stripping the gonad. For each population, sperm was collected individually for each male while oocytes of all females were pooled. Eggs were sieved on a 20 µm and 100 µm screens to remove small and large debris, respectively, the eggs being retained on the 20 µm screen. Then, the pool of eggs was divided by the number of males, and each subgroup was fertilized by a male. Fifteen minutes after fertilization, all subgroups were mixed, and all fertilized and unfertilized eggs were placed in fourteen 30 L tanks at a density of 34 to 100 eggs per mL (**Supplementary File 1, Table S2**). Thus, depending on the population, between one to three million eggs were added into each 30 L conical tank. Tanks were in a batch system containing 26 °C filtered and UV-treated seawater, supplemented with gentle air-bubbling. Larval farming density were 10 larvae per ml at day 2, and 3 larvae per ml at day 7. Seawater was changed three times per week, and larvae were fed daily with *Isochrysis galbana*, supplemented with *Skeletonema costatum* from day 7.

### Exposure of oyster larvae with microorganisms

For each population, seven conditions were tested, each using two 30 L replicate tanks. Larvae were either unexposed or exposed to microbiota from donor oysters (ME seawater D0-D14) or to the five different multi-strains bacterial mixes at two different larval developmental window (Brest D0-D14, Brest D7-D14, La Tremblade D0-D14, La Tremblade D7-D14, Arcachon D0-D14, Thau D0-D14 and Multi-site D0-D14, Multi-site D7-D14 (**Figure 1**). For ME seawater D0-D14, larvae were exposed to the whole natural microbiota coming from healthy donor oysters (Microorganism-Enriched seawater = ME seawater). This microorganism community was introduced thanks to donor oysters of microbiota which were placed into the rearing tanks. Oyster donors of microbiota were NSI (Naissains Standardisés Ifremer, standardised Ifremer spats) (Petton et al. 2013, 2015) which were always kept in controlled facilities using UV-treated seawater, strict biosecurity zoning and management procedures. The oysters were tested negative for the three main pathogens (*Vibrio coralliilyticus,* OsHV-1 µVar and *Haplosporidium costale*) of *C. gigas* from larvae to juveniles (Azéma et al. 2017; Dégremont et al. 2021). The microorganisms were added to the larvae either 3 hours post-fertilization (pf) and at each water change until day 14 pf or from day 7 pf to day 14 pf (**Figure 1**). The water changes at day 14 was performed without addition of the bacterial mixes. In this sense, the microbial exposure ended up at day 14.

**Figure 1:**
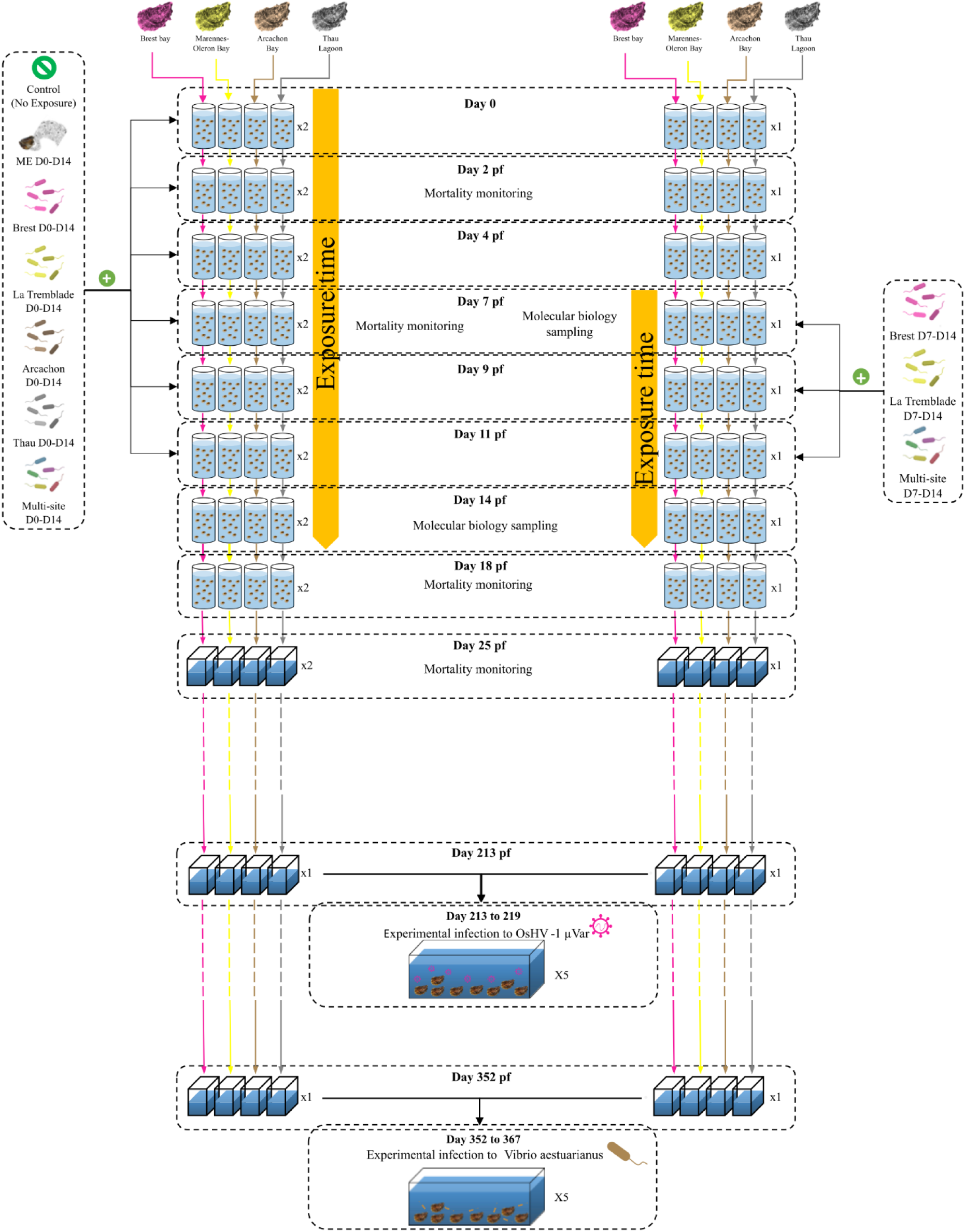
Overall experimental design for larval microbial exposure and experimental infections. Multi-parental reproduction was performed for the four oyster populations and the larvae were placed in 30 L tanks in a batch system containing 26°C filtered and UV-treated seawater, supplemented with gentle air-bubbling. Three hours post-fertilisation (pf), larvae remained unexposed (2 tanks) or were exposed in duplicate to microbiota from donor oysters (ME seawater D0-D14) or to the five different multi-strains bacterial mixes (Brest D0-D14, La Tremblade D0-D14, Arcachon D0-D14, Thau D0-D14 and Multi-site D0-D14). This microorganism exposure was renewed three times per week and lasted for 14 days. In parallel, exposure to three multi-strains bacterial mixes (Brest D7-D14, La Tremblade D7-D14 and Multi-site D7-D14) was performed on older larvae between D7 and D14 pf. During the larval stage, seawater and larvae were sampled at days 2, 7, 11, 14, 18 and 25 pf to perform growth and mortality monitoring, or to perform molecular analysis. After the larval stage, spat grew in our controlled facility. At day 213 pf (approximatively seven months old), a first set was used to carry out an experimental infection to OsHV-1 µVar and at day 352 pf (approximatively one year), a second set was used to perform a *V. Aestuarianus* experimental infection.

Larval survival was determined by counting the larvae either at days 2, 7 and 18 for oysters exposed from day 0 pf to day 14 pf or at day 18 for oysters exposed from day 7 pf to day 14 pf. Fixation rate was determined at day 25 pf for all conditions. Larvae (Pools of 10000-20000 individuals) were sampled either at days 7 pf or at day 14 pf, flash frozen in liquid nitrogen and stored at –80°C for subsequent molecular analysis. After the rearing steps, only one replicate was kept to perform the experimental infections.

All oyster populations were kept in controlled facilities of the La Tremblade hatchery using UV-treated seawater until experimental infections by OsHV-1 µVar or *V. aestuarianus*.

### OsHV-1 µVar experimental infection by cohabitation

OsHV-1 µVar experimental infection was performed either on control or microorganisms exposed oysters (seven-month-old, mean individual weight = 2.80 ± 0.69g). A randomized complete block design composed of five 50 L tanks (replicates) filled with filtered and UV-treated seawater and maintained at 20°C with adequate aeration and no food supply. Each tank contained 12 oysters of each population exposed to each condition (total: 420 oysters per tank) (**Supplementary File 2, Figure S1**). A cohabitation protocol, adapted from (Schikorski et al. 2011) was used as described. This approach starts with the injection of 100 µL of OsHV-1 µVar suspension (10^5^ OsHV-1 µVar genomic units) into the adductor muscle of pathogen-free oysters donors. This protocol allows for pathogen transmission through the natural infectious route to oysters of interest (recipient oysters). The OsHV-1 µVar donor oyster pool was composed of 25% of F15 family oysters, 25% of F14 family oysters which are POMS susceptible oysters (de Lorgeril et al. 2018) and 50% of genetically diversified NSI oysters (∼50% of susceptibility). The ratio was 1 donor oyster for 1 recipient oyster. Immediately after OsHV-1 µVar injection into donors (adductor muscle), recipient and donor oysters were uniformly distributed in each of the five experimental tanks. After 48 hours of cohabitation, all donor oysters were removed from the tanks.

In each tank, one oyster of each population exposed to each condition was sampled just before the beginning of the experimental infection (t=0h infection) and three hours post cohabitation with OsHV-1 µVar donor oysters (t=3h infection) to perform molecular analysis on whole tissue samples. The shell was removed, the whole flesh flash frozen into liquid nitrogen and stored at –80°C until it was grounded in liquid nitrogen (Retsch MM400 mill) to a powder that was then stored at –80°C until DNA and RNA extraction.

The mortality was recorded daily during eight days. Dead recipient oysters were removed daily from the tanks. During the mortality monitoring, 1 mL of water in each tank was sampled every day for the detection and the quantification of OsHV-1 µVar.

### *Vibrio aestuarianus* experimental infection by cohabitation

*Vibrio aestuarianus* experimental infection was performed either on control or microorganisms exposed oysters (12 months old; mean individual weight = 9.42 ± 1.29g) with a cohabitation protocol previously described in (Azéma et al. 2017). A randomized complete block design composed of five 100L replicate tanks filled with filtered and UV-treated seawater and maintained at 20°C with adequate aeration and without food were used. Each tank contained 10 oysters of each population exposed to each condition (total: 350 oysters per tank). The *V. aestuarianus* 02/041 strain (Garnier et al. 2008) was grown in Zobell medium at 22°C for 24h under agitation. The bacterial concentration was determined by spectrophotometry at 600nm and adjusted to an optical density (OD_600_) of 1 representing 5.10^8^ bacteria per mL. *V. aestuarianus* donor oysters were injected in the adductor muscle with 100µL of the *V. aestuarianus* 02/041 suspension and were then equally distributed among the five tanks. The *V. aestuarianus* donor oyster population was composed of an equi-number of the four oyster populations produced for this project (Brest, La Tremblade, Arcachon and Thau populations). Immediately after *V. aestuarianus* injection into donors, donor oysters were added to the five tanks containing the recipient oysters. A ratio of 1 *V. aestuarianus* donor oyster for 1.5 recipient oyster was used. After 48 hours of cohabitation, *V. aestuarianus* donor oysters were removed from the tanks.

The mortality was recorded daily during 15 days, and all the dead oysters were removed from the tanks. During the mortality monitoring, 1 mL of water in each tank was sampled every day for the detection and the quantification of *V. aestuarianus*.

### Statistical Analysis of oyster mortality

Oyster mortality rates were compared between the different microorganisms exposure set using survival analysis performed on R (v 4.2.1) (R Core Team 2022) with the package survminer (v 0.4.9) (https://cran.r-project.org/web/packages/survminer/index.html). The Kaplan-Meier method was used to represent the cumulative survival rate and log-rank test to determine the difference between conditions. A multivariate Cox proportional hazards regression model was used to compute Hazard-Ratio (HR) with confidence intervals of 95%.

### Oysters and water Genomic DNA extraction and sequencing

DNA extraction from larvae (pool of 10000 to 20000 individuals) collected during microorganisms exposure was extracted with the DNA from the tissue Macherey-Nagel kit according to the manufacturer’s protocol. Prior to 90 min of proteinase K lysis, an additional mechanical lysis was performed by vortexing samples with zirconia/silica beads (BioSpec). DNA from individual juvenile oyster tissues collected just before and during experimental infection was extracted from oyster powder with the DNA from tissue Macherey-Nagel kit according to the manufacturer’s protocol. Prior to 90 min of proteinase K lysis, an additional 12-min mechanical lysis (Retsch MM400 mill) was performed with zirconia/silica beads (BioSpec). DNA extraction from water collected during microorganisms exposure and experimental infections was extracted with the DNA from tissue Macherey-Nagel tissue kit following the manufacturer support protocol for genomic DNA and viral DNA from blood sample.

DNA concentration and purity were checked with a Nanodrop ND-1000 spectrometer (Thermo Scientific).

### qPCR analysis

Detection and quantification of OsHV-1 µVar and *V. aestuarianus* was performed by real-time quantitative PCR. All amplification reactions were performed on Roche LightCycler® 480 Real-Time thermocycler. Each reaction was carried out in triplicate in a total volume of 10 µL containing the DNA sample (2.5 µL), 5 µL of Takyon™ SYBER MasterMix blue dTTP (Eurogentec, ref UF-NSMT-B0701) and 1 µL at 500 nM of each primers for OsHV-1 µVar (OsHVDPFor5’-ATTGATGATGTGGATAATCTGTG and OsHVDPFor 5’-GGTAAATACCATTGGTCTTGTTCC) (Webb et al. 2007) and for *V. aestuarianus* (DNAj-F 5′-GTATGAAATTTTAACTGACCCACAA and DNAj-R 5′-CAATTTCTTTCGAACAACCAC) (Saulnier et al. 2009). qPCR cycling conditions were as follows: 3 min at 95°C, followed by 45 cycles of amplification at 95°C for 10 s, 60°C for 20 s, and 72°C for 30s. After these PCR cycles a melting temperature curve of the amplicon was generated to verify the specificity of the amplification. The DNA polymerase catalytic subunit amplification product cloned into the pCR4-TOPO vector was used as a standard at 10-fold dilutions ranging from 10^3^ to 10^10^ copies/ml for OsHV-1 µVar quantification and genomic DNA from *V. aestuarianus* ranging from 10^2^ to 10^7^ copies/ml for *V. aestuarianus* quantification. Absolute quantification of OsHV-1 µVar or *V. aestuarianus* was calculated by comparing the observed Cp values to standard curve.

### 16S rDNA library construction and sequencing

Library construction (with primers 341F 5’-CCTAYGGGRBGCASCAG and 806R 5’-GGACTACNNGGGTATCTAAT targeting he V3-V4 region of the 16S rRNA gene) (Klindworth et al. 2013) and sequencing on a MiSeq v2 (2×250 bp) were performed by ADNid (Montpellier, France).

### RNA extraction and sequencing

RNA was extracted from oyster powder (individual) by using the Direct-Zol RNA miniprep kit (Zymo Research) according to the manufacturer’s protocol. RNA concentration and purity were checked using a Nanodrop DN-1000 spectrometer (Thermo Scientific), and their integrity was analysed by capillary electrophoresis on a BioAnalyzer 2100 (Agilent).

### RNAseq library construction and sequencing

RNA-Seq experiments were performed on 3 individuals per condition. RNA-Seq library construction and sequencing were performed by the Bio-Environment Platform (University of Perpignan, France). Stranded libraries were constructed from 500 ng of total RNA using NEBNext UltraII and sequenced on a NextSeq550 instrument (Illumina) in single-end reads of 75 bp.

### Bioinformatic pipelines for 16S rRNA gene barcoding analysis

Previously published barcoding datasets (de Lorgeril et al. 2018; King et al. 2019b; Clerissi et al. 2020, 2022; Fallet et al. 2022) from 687 POMS-resistant and 664 POMS-sensitive oysters were re-analysed in this study in order to predict bacteria which were potentially associated with oyster POMS resistant phenotypes. Datasets used for these analyses are in **Supplementary File 1, Table S3**. These datasets were individually analysed under the Toulouse Galaxy instance (https://vm-galaxy-prod.toulouse.inra.fr/) (Goecks et al. 2010) with the Find Rapidly OTU with Galaxy Solution (FROGS) pipeline (Escudié et al. 2018). In brief, paired reads were merged using FLASH (Magoč and Salzberg 2011). After denoising and primer/ adapter removal with cutadapt (Martin 2011), clustering was performed using SWARM (Mahé et al. 2014), which uses a novel clustering algorithm with a threshold (distance = 3) corresponding to the maximum number of differences between two OTUs. Chimeras were removed using VSEARCH (Rognes et al. 2016). We filtered out the data set for singletons and performed an affiliation using Blast against the Silva 16S rDNA database (release 132) to produce an OTU and affiliation tables. In order to identify bacterial taxa which were significantly overrepresented in the microbial community associated to POMS resistant compared to POMS sensitive oysters, the “LDA Effect Size” (LEfSe) method (Segata et al. 2011) was used with a normalized relative abundance matrix. This method uses a Kruskal-Wallis followed by Wilcoxon tests (pval ≤ 0.05) and then performs a linear discriminant analysis (LDA) and evaluate the effect size. The taxa with a LDA score greater than 2 were considered as significantly enriched in POMS resistant compared to sensitive oysters.

Sequencing data obtained on the samples from this study were processed with the SAMBA (v 3.0.2) workflow developed by the SeBiMER (Ifremer’s Bioinformatics Core Facility). Briefly, Amplicon Sequence Variants (ASV) were constructed with DADA2 (Callahan et al. 2016) and the QIIME2 dbOTU3 (v 2020.2) tools (Bolyen et al. 2019), then, contaminations were removed with microDecon (v 1.0.2) (McKnight et al. 2019). Taxonomic assignment of ASVs was performed using a Bayesian classifier trained with the Silva database v.138 using the QIIME feature classifier (Wang et al. 2007). Finally, community analysis and statistics were performed on R (R version 4.2.1) (R Core Team 2022) using the packages phyloseq (v 1.40.0) (McMurdie and Holmes 2013) and Vegan (v 2.6-4) (Oksanen et al. 2022). Unique and overlapping ASVs of each sample group were plotted using the UpsetR package (v 1.4.0) (Conway et al. 2017). For beta-diversity, the ASVs counts were preliminary normalized with the “rarefy_even_depth” function (rngseed = 711) from the package phyloseq (v 1.40.0)(McMurdie and Holmes 2013). Principal Coordinates Analysis (PcoA) were computed to represent dissimilarities between the samples using the Bray-Curtis distance matrix. Differences between groups were assessed by statistical analyses (Permutational Multivariate Analysis of Variance) using the adonis2 function implemented in vegan (Oksanen et al. 2022).

In order to follow the long-term installation (or not) of each of the bacteria used in the multi-strain bacterial mixes in the oyster microbiota, 16S rRNA genes sequences obtained during the identification of each of the bacteria composing the multi-strain bacterial mixes were used as a query for a similarity BLASTn search against all the ASVs sequence from the dataset (Altschul et al. 1990). A mock community composed of equal amounts of DNA from the bacteria composing the multi-strain bacterial mixes were also used as a positive control to validate our search method. ASVs sequences with a percentage of identity higher than 99% were considered present in the tested samples.

### Bioinformatic pipeline for RNA-Seq analysis

All data treatments were carried out under a local galaxy instance (http://bioinfo.univ-perp.fr) (Goecks et al. 2010). Reads quality was checked with FastQC (Babraham Bioinformatics) with default parameters (Galaxy Version 0.72). Adapters were removed using Trim Galore (Galaxy Version 0.6.3) (Babraham Bioinformatics). Reads were mapped on *C. gigas* genome (assembly cgigas_uk_roslin_v1) using RNA STAR (Galaxy Version 2.7.8a) (**Supplementary File 3: RNAseq Mapping results)** and HTSeq-count (Anders et al. 2015) was used to count the number of reads overlapping annotated genes (mode Union) (Galaxy Version 0.9.1). The differential gene expression levels were analysed with the DESeq2 R package (v 1.36.0) (Love et al. 2014). Finally, Rank-based Gene Ontology Analysis (GO_MWU package) was performed using adaptive clustering and a rank-based statistical test (Mann–Whitney U-test combined with adaptive clustering) with the following parameters: largest = 0.5; smallest = 10; clusterCutHeight = 0.25. The signed “-Log(adj pval)” (obtained from the DESeq2 analysis) was used as an input for the GO_MWU analysis. The R and Perl scripts used can be downloaded [https://github.com/z0on/GO_MWU] (Wright et al. 2015).

## Results

### 23 bacterial strains with potential beneficial effects were selected to generate the multi-strain bacterial mixes

To isolate bacteria with potential beneficial effects against oyster infectious disease, we hypothesised that bacteria should be isolated from disease resistant oysters. For this purpose, wild oysters aged between 12 and 18 months were sampled closed to farming areas. Oysters located in these areas are submitted to high pathogen pressure and have been shown to be resistant to POMS disease (Gawra et al. 2023). To maximise the biodiversity of the bacterial collection, oysters were sampled from 4 geographical French sites at two different seasons. A total of 334 bacteria were isolated (**Supplementary File 1, Table S4**); from which 166 bacteria were obtained from the February 2020 sampling campaign, and 168 bacteria from the November 2020 sampling campaign. This corresponded to 97, 144, 56, and 67 bacteria isolated from Brest, La Tremblade, Arcachon, and Thau sites, respectively. They were named according to the sampling site (“ARG” for Brest bay, “LTB” for La Tremblade in Marennes Oleron bay, “ARC” for Arcachon bay and “THAU” for Thau lagoon) followed by the number of the isolate. The 16S rRNA gene sequence was obtained for 293 strains. The identified bacteria were divided into the following phyla: *Proteobacteria* (62.8%), *Firmicutes* (15.3%), *Bacteroidetes* (12.3%) and *Actinobacteria* (9.6%) (**Figure 2**). The three major genera were *Vibrio*, *Bacillus* and *Shewanella* (**Figure 2**). The majority of the isolated species were found in all sites.

**Figure 2:**
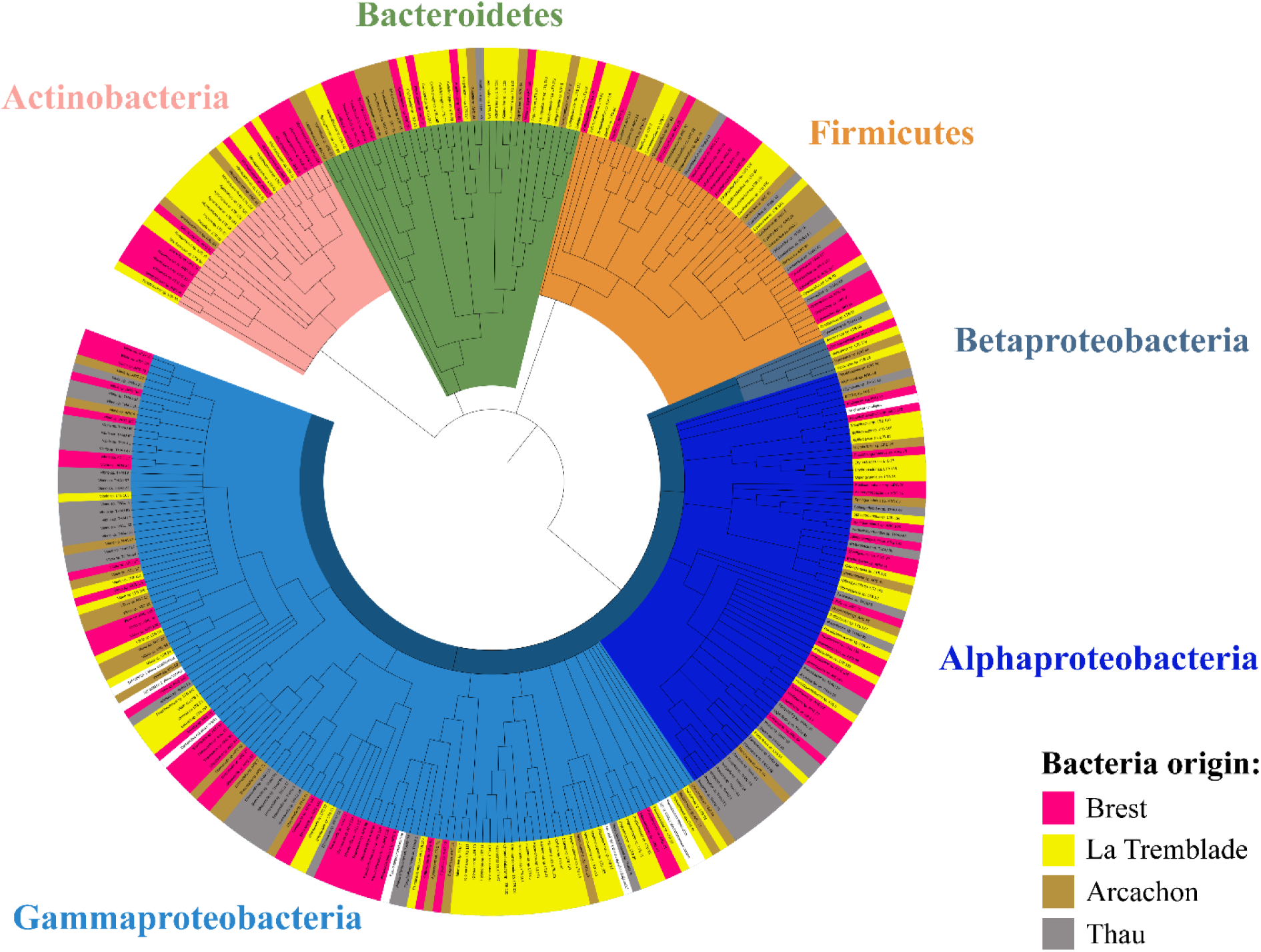
293 strains were identified in the bacterial collection sampled from POMS-resistant oysters. Phylogenetic tree of the 293 identified bacteria composing the collection of bacteria isolated from POMS-resistant oysters sampled in the Brest bay (pink), La Tremblade in Marennes-Oleron bay (yellow), the Arcachon bay (brown) and the Thau lagoon (grey) based on the V1-V5 loop alignment of bacterial 16S rDNA by a Maximum likelihood method with the Tamura-Nei parameter model in MEGA X (301 sequences) and 1000 bootstrap replicates. The collection is composed by 62.8% of Proteobacteria (different shades of blue), 15.3% of Firmicutes (orange), 12.3% of Bacteroidetes (green) and 9.6% of Actinobacteria (salmon).

In parallel, *in silico* correlation analysis was performed to predict bacteria preferentially associated with resistant or sensitive oysters. This LefSE analysis (Segata et al. 2011) was performed based on previously published 16S rRNA genes barcoding datasets which describes the bacterial part of the bacterial microbiota community isolated from 687 POMS-resistant and 664 POMS-sensitive oysters (**Supplementary File 1, Table S3**). Based on this analysis, 118 bacterial genera were shown as preferentially associated with POMS-resistant oysters (**Supplementary File 1, Table S5**). By combining the data obtained from this predictive *in silico* analysis and data from the scientific literature about bacteria shown to be beneficial in an aquaculture context (Rengpipat et al. 2000; Zhang et al. 2009; Kesarcodi-Watson et al. 2012; Touraki et al. 2012; Sun et al. 2013; Guzmán-Villanueva et al. 2014; Yan et al. 2014; Reda and Selim 2015; Tan et al. 2016; Chauhan et al. 2017; Makled et al. 2017; Lv et al. 2019), we selected 12, 17, 10 and 8 bacteria for the Brest, La Tremblade, Arcachon and Thau sites respectively (**Table 1**). These bacterial strains were then tested for their cytotoxic effects on 2 days old larvae. The most cytotoxic bacteria were discarded. Based on these results, we kept five, seven, five and five site-specific bacteria to produce the Brest, La Tremblade, Arcachon and Thau multi-strain bacterial mixes respectively (**Table 1**). A fifth multi-site bacterial mix was produced from bacteria isolated from oysters sampled on all sites. For this purpose, seven different bacteria were chosen because they displayed the least cytotoxic effects on larvae (**Table 1**).

In summary, we collected bacteria from disease-resistant oysters. We then combined our findings with existing literature and utilized *in silico* predictive analysis. This allowed us to create four site-specific and one multi-site multi-strain bacterial mixes, all of which have the potential to benefit oyster health.

### Microorganisms exposure during larval rearing induces long term protection against POMS and Vibriosis which relies on bacterial mix composition and oyster origin

The multi-strain bacterial mixes were added to four oyster populations during the larval rearing. The four populations were the sympatric oysters from which the bacteria were isolated (*i.e.,* Brest, La Tremblade, Arcachon and Thau). An exposure with a whole microbiota community coming from healthy hatchery donor oysters was also performed (ME seawater D0-D14). This oysters were shown to be devoid of the three main pathogens (*V. coralliilyticus,* OsHV-1 µVar and *Haplosporidium costale*) of *C. gigas* from larvae to juveniles (Azéma et al. 2017; Dégremont et al. 2021). Oyster’s larvae were exposed to bacterial mixes either from blastula (3h post-fertilization (pf)) to pediveliger stage (14 days pf) (D0 to D14) or from veliger stage (seven days pf) to pediveliger stage (14 days pf) (D7 to D14) (**Figure 1**). Overall, these microorganisms exposures during larval rearing stages displayed from moderate to strong effect on larval survival. These effects rely on oyster origins and, also, on the bacterial content of the microorganism exposure (**Supplementary File 4 Effect of bacterial mixes on oyster larvae and Supplementary File 2, Figure S2)**.

Subsequently, each oyster population (exposed and control) were challenged with OsHV-1 µVar infection during juvenile stages or *V. aestuarianus* during adult stages. The success of the experimental infection was verified by quantifying the viral or Vibrio DNA concentration in the sea water of the experimental tanks (**Supplementary File 1, Table S6 and Table S7**).

In response to OsHV-1 µVar infection, a significant reduction of the mortality risk of 21% (Log-rank test: pval = 0.038), 25% (Log-rank test: pval = 0.009), and 28% (Log-rank test: pval = 0.008) was observed in the oysters (all populations combined) exposed to the Arcachon D0-D14, La Tremblade D7-D14 and D0-D14 ME seawater mixes, respectively (**Figure 3**). We observed that the mortality started 3 days after the POMS disease induction, and differences between the control and exposed samples appeared as soon as mortality started for oysters exposed to the Arcachon D0-D14, La Tremblade D7-D14 and, ME seawater D0-D14 oysters (**Supplementary File 2, Figure S3**).

**Figure 3:**
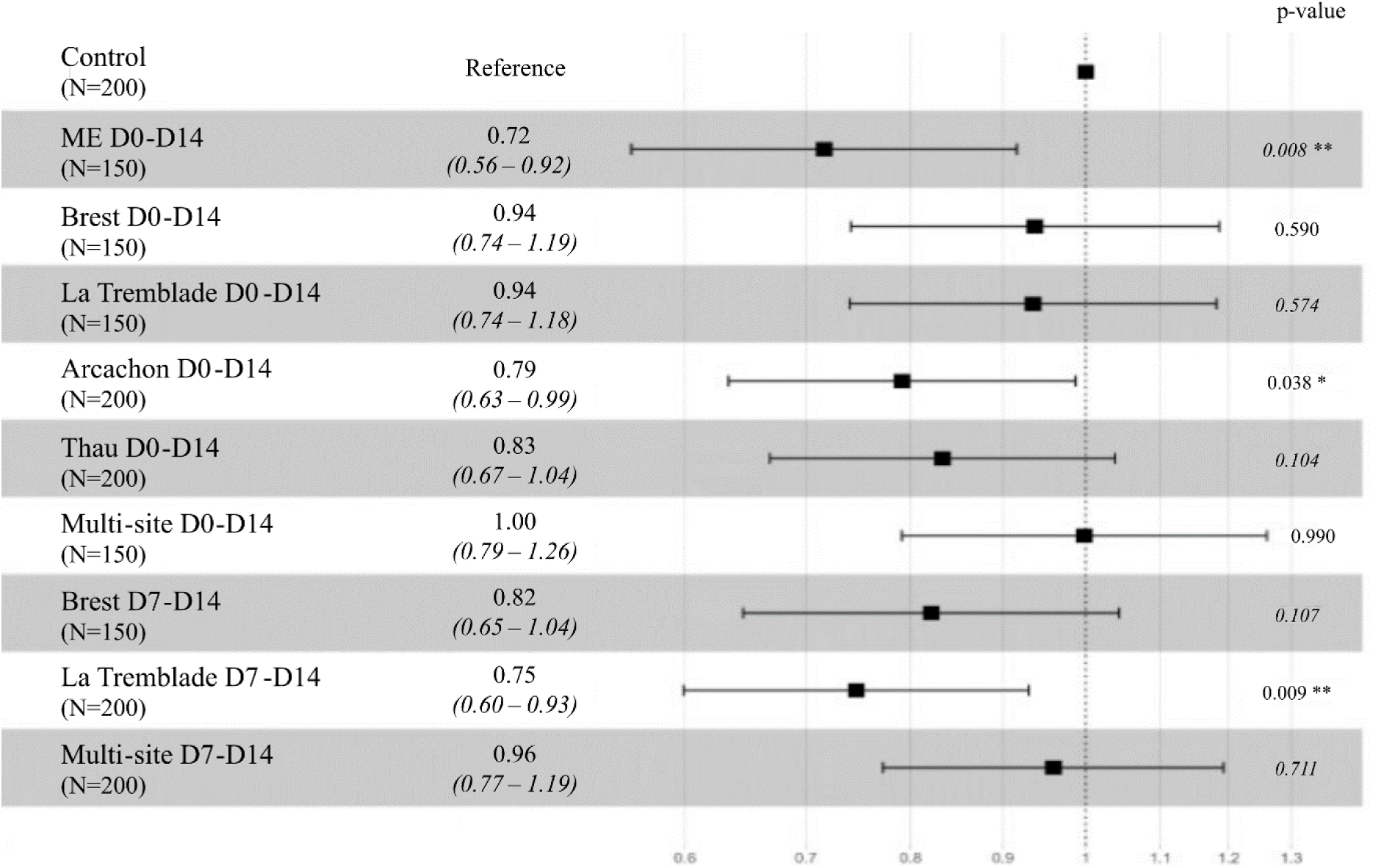
Bacterial mixes and ME-seawater exposure during larval rearing reduce the mortality risk induced by POMS. Forest plot representing the Hazard-Ratio value of mortality risk during the OsHV-1 µVar experimental infection for oysters (all populations combined) exposed to microorganisms compared to control oysters. The numbers into brackets under the different conditions correspond to the number of oysters used during the experimental infection. The Hazard-Ratio value is indicated to the right of the conditions, except for the control condition, which is indicated as reference. The p-value of the log rank test is indicated on the right-hand side of each row.

In response to vibriosis, a significant reduction of the mortality risk of 28% (Log-rank test: pval = 0.006) was observed for the ME seawater D0-D14 exposed oysters (**Figure 4**) (**Supplementary File 2, Figure S4**). Other exposures did not lead to reduction of mortality.

**Figure 4:**
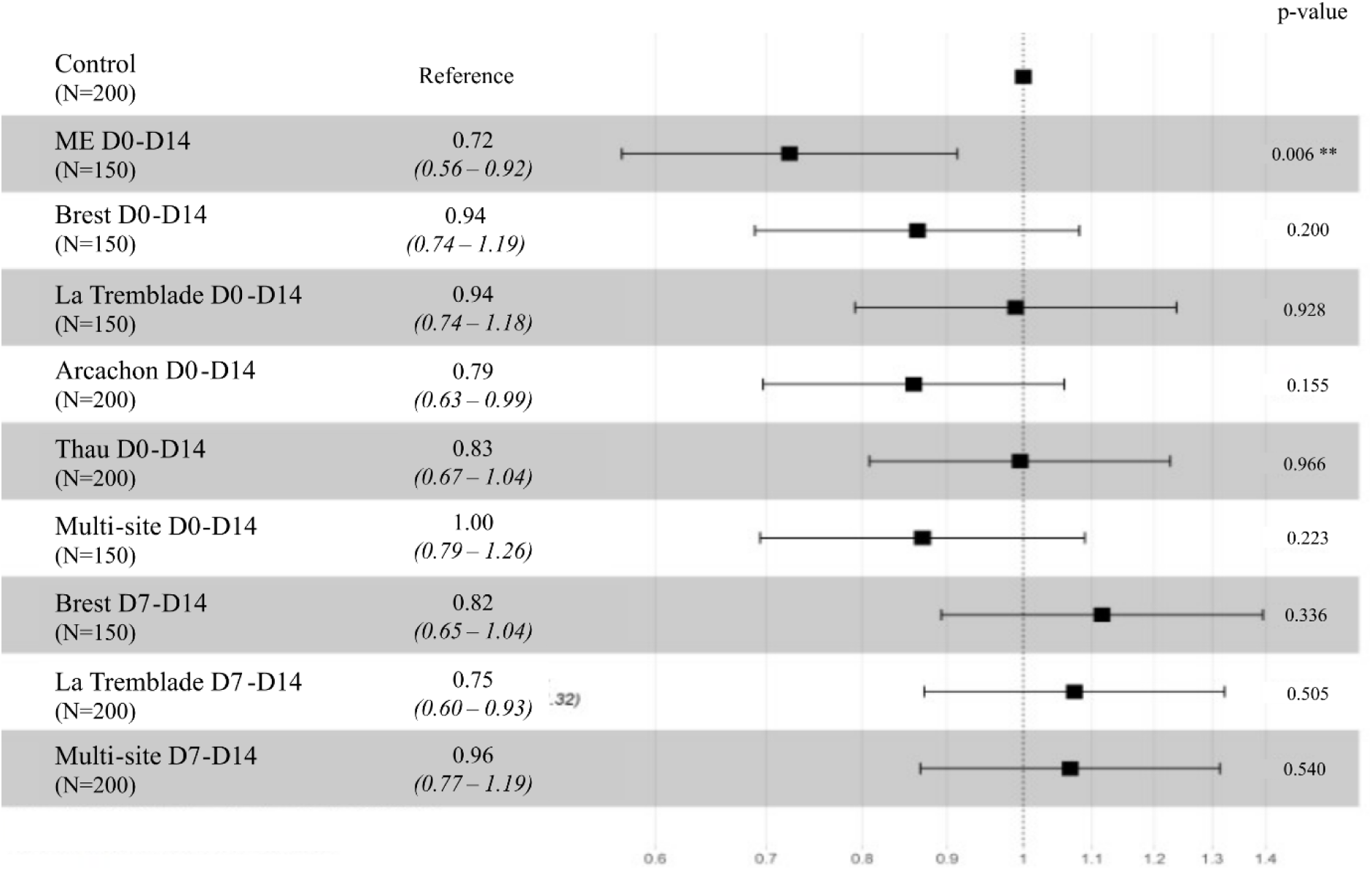
ME-seawater exposure during oyster larval rearing can reduce the mortality risk induced by *V. aestuarianus*. Forest plot representing the Hazard-Ratio value of mortality risk during the *V. aestuarianus* experimental infection for oysters (All populations confounded) exposed to microorganisms compared to control oysters. The numbers into brackets under the different conditions correspond to the number of oysters used during the experimental infection. The Hazard-Ratio value is indicated to the right of the conditions, except for the control condition, which is indicated as reference. The p-value is indicated on the right-hand side of each row.

For both Vibriosis and viral infection, the beneficial effect in response to each of the mixes depended on the oyster origin (**Supplementary File 2, Figure S3)**. Oysters originating from Arcachon showed the best reduction in mortality in response to both infections regardless of the bacterial exposure conditions during the larval stages. The effect of the microorganisms exposure was intermediate on oysters from La Tremblade and less pronounced on oysters from Brest and Thau (**Supplementary File 2, Figure S3**).

In summary, larval exposure to bacterial mixes or Microorganism-Enriched seawater (ME seawater) conferred a beneficial effect on the survival of the oysters against the POMS disease in juvenile oysters while only Microorganism-Enriched seawater (ME seawater) conferred a beneficial effect against Vibriosis. No preferential beneficial effect was nevertheless observed when the oysters were exposed to their sympatric compared to allopatric strains.

### Microorganism exposure during larval rearing induced long term changes of the microbiota composition

To test the immediate and long-term effect of the microorganism exposure on the oyster microbiota composition, we analysed the bacterial communities by 16S rRNA gene sequencing during the larval stage after seven days of exposure and during the juvenile stage seven months after the exposure. We focused our study on the three conditions of bacterial exposure that conferred significant increase on the survival of oysters during OsHV-1 µVar and *V. aestuarianus* experimental infection.

Sequencing of the V3-V4 hypervariable region of the 16S rRNA gene resulted in a total of 10,868,202 clusters. After quality check (deleting primers and low-quality sequences, merging, and removing chimeras) and ASV clustering, 5,322,399 reads (49%) with an average of 35,962 reads per sample were retained for downstream analyses.

A higher species richness was observed seven days after the exposure for ME-seawater exposed larvae but not after exposure to bacterial mixes (**Figure 5 A,B**). This difference was not maintained at juvenile stage **(Figure 5 C)**. Dissimilarity analysis, based on the Bray-Curtis index, showed that the larvae microbiota composition differed between conditions after seven days of microorganism exposure, whatever the condition (**Table 2)**. This difference remained statistically significant at juvenile stage for ME seawater D0-D14 and La Tremblade D7-D14 conditions (**Table 2)**.

**Figure 5:**
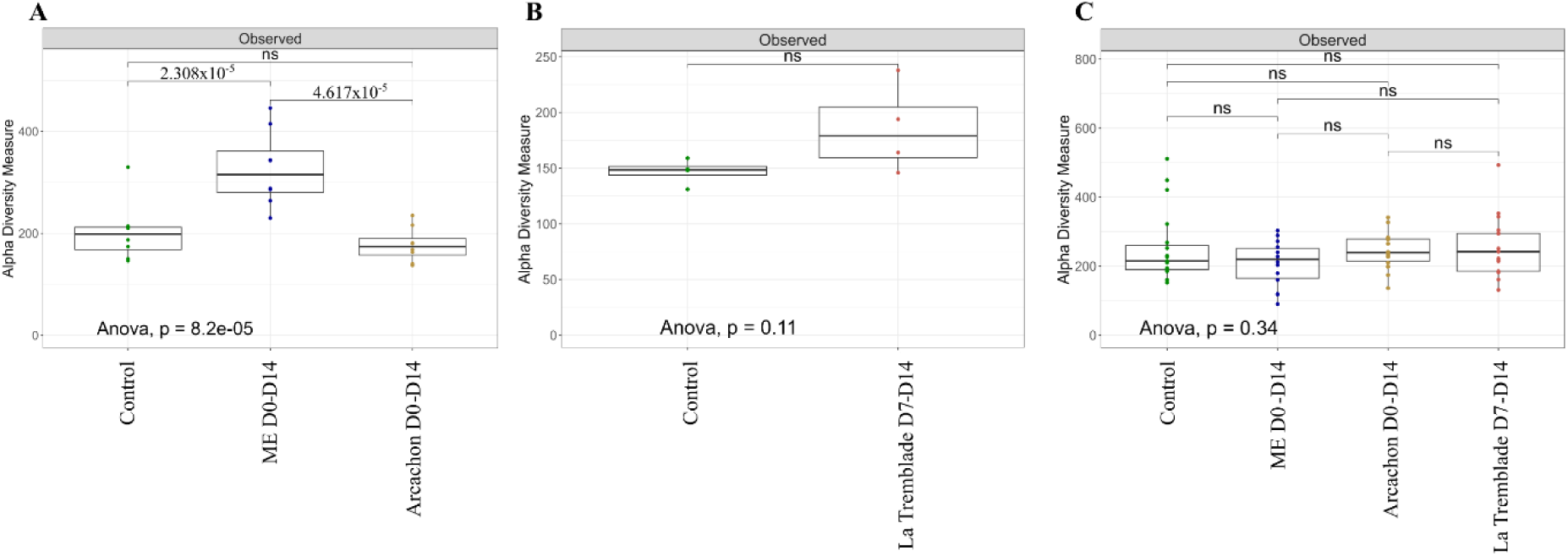
The richness of oyster microbiota is transiently increased after larval exposure to Microorganism-Enriched seawater. The alpha-diversity indexes (observed species richness) of larvae microbiota after seven days of exposure (A,B), or juvenile microbiota seven months after the exposure (C) are indicated. For larval stages (A) and (B), analyses were performed on all oyster populations confounded which represent eight pools of 10000-20000 larvae sampled in eight independent tanks for exposure to ME D0-D14 and Arcachon D0-D14 (A) and on four pools of 10000-20000 larvae sampled in four independent tanks for exposure to La Tremblade D7-D14 (B). For juvenile stages (C), analyses were performed on all oyster population confounded which represent 68 individuals sampled in five independent tanks. Significant changes are indicated by their p-value and “ns” stands for “not significant”.

**Table 2:** Long-lasting modifications in *C. gigas* microbiota composition occurred following microorganisms exposure. Permanova (adonis2) on the Bray-Curtiss dissimilarity matrix showing the effects of microbial exposure on microbiota community compared to control condition for larvae after seven days of microbial exposure and for juveniles seven months after the microbial exposure. For larvae, analyses were performed on all oyster populations confounded which represent eight pools of 10000-20000 larvae sampled in eight independent tanks for exposure to ME D0-D14 and Arcachon D0-D14 and on four pools of 10000-20000 larvae sampled in four independent tanks for exposure to La Tremblade D7-D14. For juvenile stages, analyses were performed on all oyster population confounded which represent 68 individuals sampled in five independent tanks.

| Conditions<br>(Compared to<br>control) | Larvae (after 7 days of exposure) |  |  |  | Juvenile (7 months) |  |  |  |
| --- | --- | --- | --- | --- | --- | --- | --- | --- |
|  | Dum Sq | R<sup>2</sup> | F | p | Dum Sq | R<sup>2</sup> | F | p |
| ME D0-D14 | 0.75 | 0.18 | 4.46 | 0.001 | 0.19 | 0.04 | 1.45 | 0.026 |
| Arcachon D0-D14 | 0.85 | 0.19 | 3.30 | 0.001 | 0.08 | 0.02 | 0.75 | 0.908 |
| La Tremblade D7-D14 | 0.43 | 0.29 | 2.98 | 0.036 | 0.17 | 0.04 | 1.55 | 0.033 |

We additionally checked for the presence of the added bacteria, during the larval stage, after seven days of exposure with the last addition of bacteria done 48 hours before sampling, and at juvenile stage seven months post-exposure. Two bacterial strains out of the 5 added in larvae exposed to Arcachon D0-D14 were retrieved and represented 3.3 to 25.9 % of the total bacterial community (**Supplementary File 2, Figure S5A**). ASVs associated with the added bacteria of the La Tremblade D7-D14 ranged from 0.09 to 0.96 % in the corresponding larvae samples (**Supplementary File 2, Figure S5C**). None of the ASVs corresponding to bacteria used for the exposure could be detected at the juvenile stages seven months post-exposure (**Supplementary File 2, Figure S5B,D)**. Furthermore, either for larvae or juvenile oysters, bacterial strains did not show a preference for implantation in their sympatric host population (**Supplementary File 2, Figure S5**). Using this pipeline of detection, we were able to detect these ASVs on a mock control containing an artificial mix of bacteria in the same proportion except for *Paracococcus sp.* LTB95 and *Psychrobacter sp.* LTB83 (**Supplementary File 2, Figure S6)**. This indicated that the lack of detection of the ASVs in exposed oyster is due to an absence of the bacteria rather than a technical shortcoming in our detection pipeline, except for *Paracococcus sp.* LTB95 and *Psychrobacter sp.* LTB83.

In summary, a few proportions of the different bacteria that were added during the larval rearing were detected in the oyster microbiota 48h after the last addition of bacteria, and none of them were maintained on a long-term basis. Despite this lack of bacterial colonization, the overall composition of the microbiota was modified in response to the bacterial exposure and these changes remained up to the juvenile stages.

### Microorganisms exposure during larval rearing induced long-term changes in oyster immunity

The long-term impact of the microorganisms exposure on oyster gene expression was analysed by RNA-seq on juvenile oysters before and during POMS challenge. In total, RNA sequencing produces between 15.1 and 36.6 million reads per sample (mean number of reads = 26 millions). Among these reads, 67.28% to 77.52% were mapped on *C. gigas* reference genome (assembly cgigas_uk_roslin_v1) (**Supplementary File 3: RNAseq Mapping result)**.

For each of the four oyster populations, the number of differentially expressed genes (DEGs) in oysters exposed to bacterial mixes or to ME seawater compared to control oysters, was higher before the infection than 3h after the beginning of the infection except for the condition where Brest oysters were exposed to ME seawater (**Figure 6**). Furthermore, each oyster population displayed a specific transcriptomic response, which strongly varied according to the microorganism exposure. (**Figure 7**).

**Figure 6:**
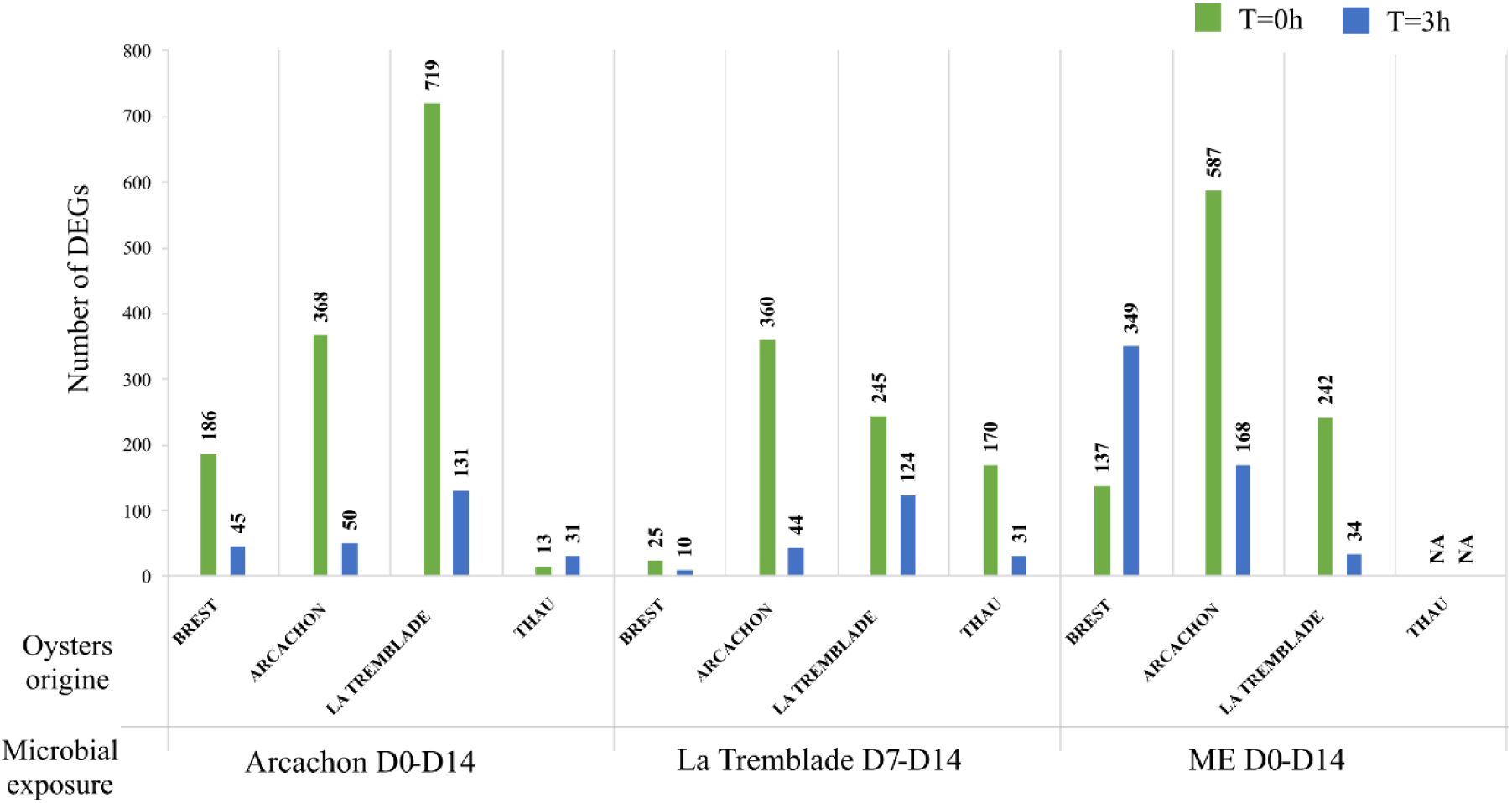
Long-lasting changes in gene expression was observed in juvenile oysters seven months after larval exposure. Histogram of differentially expressed genes (DEGs) in oysters exposed to Arcachon D0-D14, La Tremblade D7-D14 or ME seawater D0-D14 compared to control oysters for the four oyster populations (Brest, Arcachon, La Tremblade and Thau) prior to OsHV-1 µVar infection (green) and 3h post infection (blue). n=3 individuals per condition.

**Figure 7:**
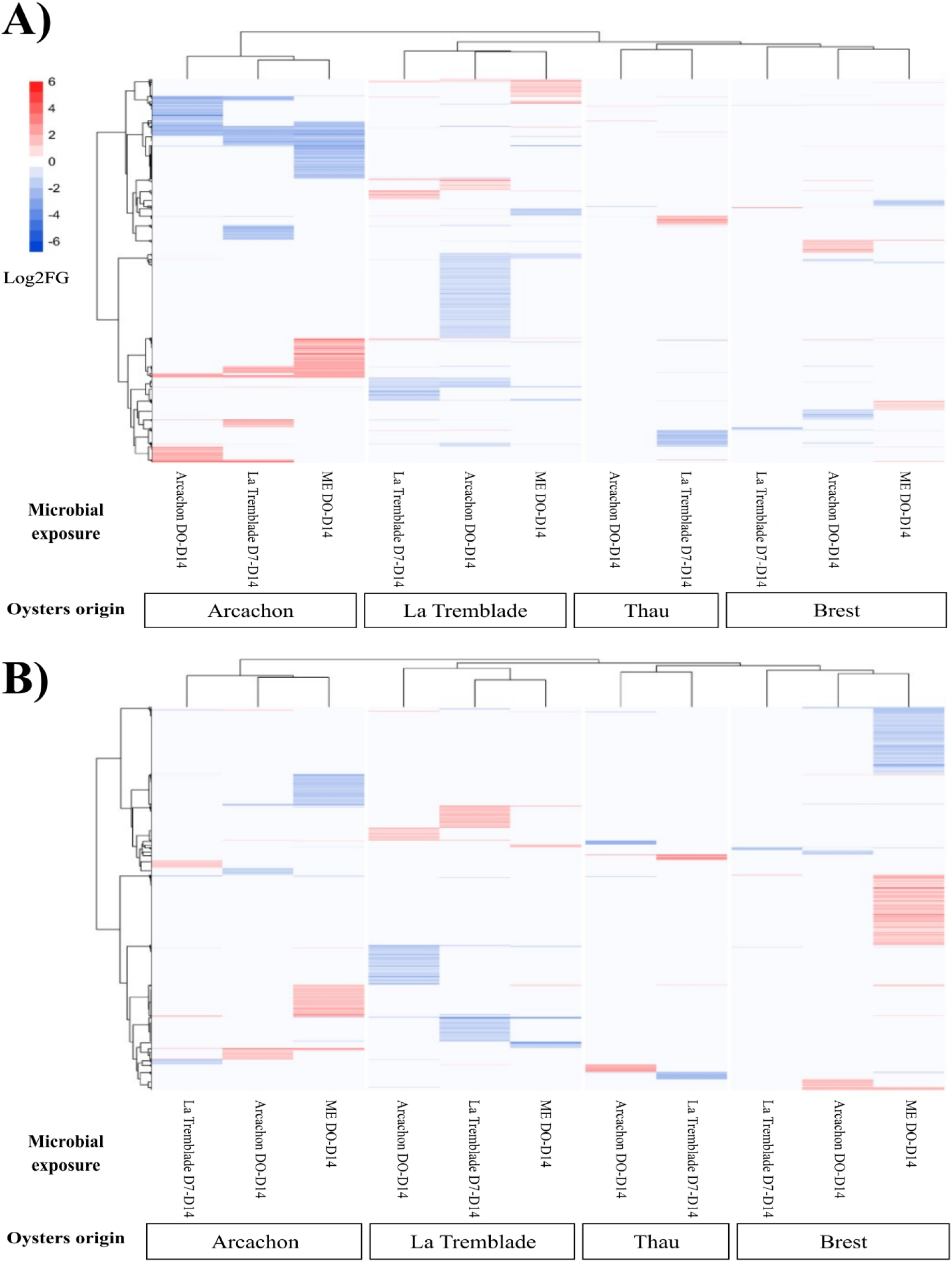
Specific gene expression profiles were observed in response to each microorganism exposure. Heatmap of differentially expressed genes (DEGs) in oysters exposed to Arcachon D0-D14, La Tremblade D7-D14 or ME seawater D0-D14 compared to control oysters for the four oyster populations (Brest, Arcachon, La Tremblade and Thau) (A) prior to OsHV-1 µVar infection and (B) 3h post OsHV-1 µVar infection. The intensity of DEG ratios is represented by the Log2 Fold Changes (Log2FC) for over expressed DEGs (in red) and under expressed DEGs (in blue). n=3 individuals per condition.

To identify which biological processes were affected by the microbial exposure, we conducted a Rang-Based Gene Ontology Analysis (GO_MWU) (Wright et al. 2015). The range of biological process enriched in DEGs (microorganisms exposed vs control) before and during the onset of the POMS disease included many GO terms such as, metabolism, RNA and DNA process, protein processing, signal transduction, transport, and immune functions. We then focused on the enriched immune functions in oysters exposed to microorganisms compared to the control oysters (**Figure 8**). The most significantly enriched functions related to immunity across all oyster populations and all treatments were general functions of immunity (defence response, immune system process), functions related to the response to organisms (response to bacterium, response to virus), a function related to the positive regulation of response to stimulus and a function related to G-protein signalling pathway (**Figure 8**). As the oysters from Arcachon showed the greatest reduction in mortality risk in the face of viral infection and *V. aestuarianus*, with all the microbial exposures, we then analysed, for these oysters only, the individual DEGs for the main enriched functions linked to immunity described in (**Figure 8**). This analysis revealed that before the infection (t=0), gene coding for Pattern Recognition Receptor (PRRs) (C-type lectins, C1q domain containing protein), innate immune pathways (toll-interleukin receptor (TIR), Complement pathway), interaction with bacteria (Bactericidal permeability-increasing protein) and antiviral pathways (RNA and DNA Helicases, RNA-dependent RNA polymerase) were found to be over-represented in microbial exposed oysters compared to control oysters (**Figure 9**) (**Supplementary File 5 List of DEGs**).

**Figure 8:**
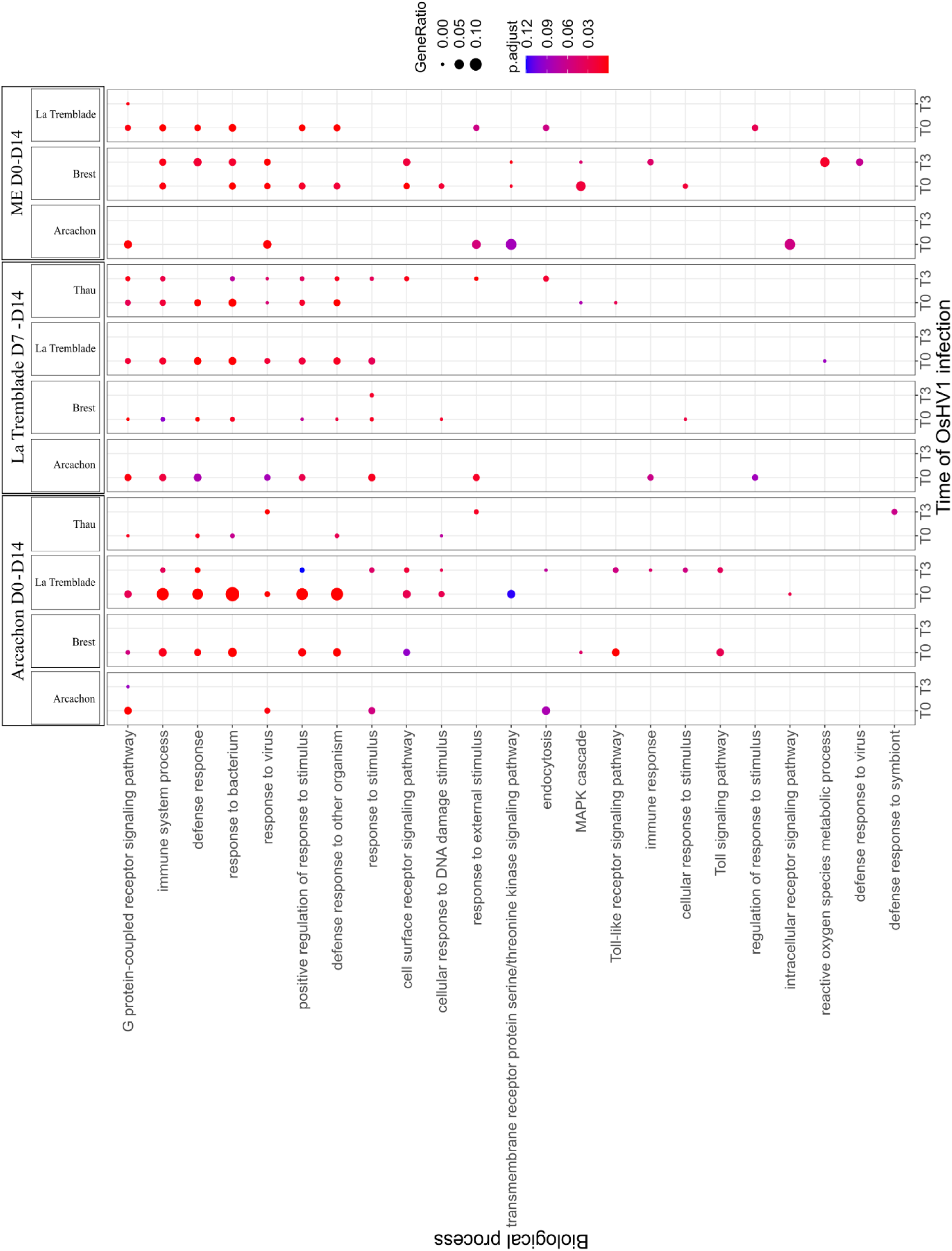
GO term enrichment analysis revealed important immune pathways modified in response to the microorganism exposure. Dot plot showing the overrepresented GO terms (FDR <0.1) of biological process (BP) related to immune function identified using GO_MWU for the four oyster populations (Brest, Arcachon, La Tremblade and Thau) exposed to Arcachon D0-D14, La Tremblade D7-D14 or ME seawater D0-D14 compared to control oysters at t=0 and t=3h OsHV-1 µVar infection. The dot size is proportional to the number of differentially expressed genes (DEG) in the biological process compared to the control condition, and the colour of the dot shows the significance.

**Figure 9:**
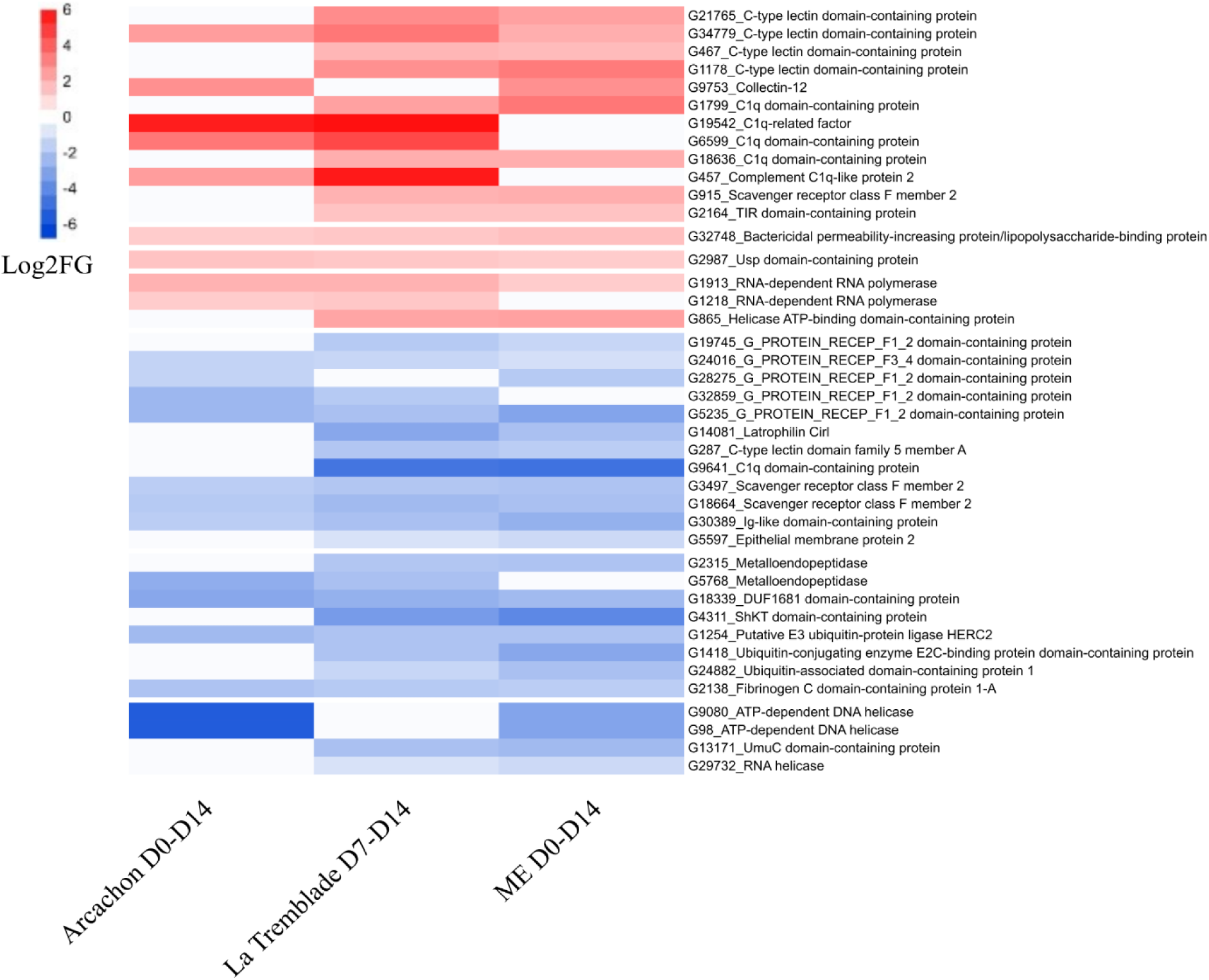
Detailed immune-related gene expression revealed key genes modified in Arcachon oysters in response to microorganism exposure. Transcriptomic response of immune related genes for oysters of the Arcachon population exposed to Arcachon D0-D14, La Tremblade D7-D14 or ME seawater D0-D14 compared to control condition before OsHV-1 µVar experimental infection. Heatmap of DEGs associated with immune processes. Only DEGs found under at least two conditions of exposure to micro-organisms were shown. The intensity of DEG ratios is expressed in Log2 Fold changes (Log2FC) for over expressed DEGs (in red) and under expressed DEGs (in blue).

In summary, long-lasting changes in gene expression were observed in juvenile oysters seven months after they had been exposed to bacterial mixes or Microorganisms Enriched seawater during larval stages. The long-lasting transcriptional responsiveness was found to be influenced by the host’s origin, was specific to the type of treatment and significantly impacts the host immune response.

## Discussion

OsHV-1 µVar, a threatening pathogen for oyster production, has spread not only in Europe (Segarra et al. 2010; Petton et al. 2021) but also to the United States (Friedman et al. 2005), Japan (Shimahara et al. 2012), Australia (Paul-Pont et al. 2013), China (Bai et al. 2015) and New-Zealand (Delisle et al. 2022). On the other hand, the pathogenic bacterium *V. aestuarianus* has been observed to spread across Europe (Mesnil et al. 2022). Innovative research and concerted efforts are currently being explored for safeguarding *C. gigas* and ensuring the sustainability of oyster farming on a global scale (Green and Montagnani 2013; Dégremont et al. 2015, 2020; Lafont et al. 2017). One promising avenue of research involves education of the oyster immune system through proper setting of the microbiota during early life. Similar to the way early microbial colonization impacts human health (Gensollen et al. 2016; Renz et al. 2017), introducing specific microorganisms to oyster larvae can potentially educate their innate immune systems and improve disease resistance (Galindo-Villegas et al. 2012; Fallet et al. 2022). The immune system in oysters is set up early during the development since the existence of a primitive immune system has been detected in the trochophore larva (Tirapé et al. 2007; Liu et al. 2015). This microbial education plan is a promising strategy as it is easy to implement, not costly and, can be performed on numerous animals (several hundred million of larvae) at the same time by bath or on their diet. However, a challenge arises in the form of current hatchery practices, which aim to minimize the introduction of both non-pathogenic and pathogenic microorganisms into larval tanks (Bourne et al. 1989; Helm et al. 2004; Eljaddi et al. 2021; Cordier et al. 2021). Mortality issues, particularly during larval rearing, have led to the use of antibiotics in hatcheries. Therefore, finding a balance between educating the immune system and addressing concerns about uncontrolled microbiota transfer is crucial. Here, our study explored the feasibility of microbial education in oyster larvae while considering and mitigating the risks associated with uncontrolled transfer of hazardous microorganisms.

For this purpose, we investigated the long-term protection conferred by a larval exposure to a controlled non-pathogenic whole microbiota transferred from donor oysters. The donor oysters used in this study were always kept in biosecured facilities. In this way, the oysters were shown to be devoid of the three main pathogens of *C. gigas* from larvae to juveniles (Azéma et al. 2017; Dégremont et al. 2021). In parallel, we performed the same assay using a reduced, synthetic bacterial community composed of cultivable bacteria. The cultivable bacteria were isolated from POMS-resistant oysters and selected according to their predictive beneficial effect on POMS disease based on robust correlation analysis. The microbial exposures were performed on 4 different oyster populations each exposed to either a sympatric or allopatric multi-strain bacterial mixes. We showed that larval exposure to a whole microbiota from donor oysters provided protection against both the POMS disease and *V. aestuarianus* infection. In a different way, larvae exposed to multi-strains bacterial mixes showed improved survival against OsHV-1 µVar but no protection against *V. aestuarianus* infection. The host origin was identified as a critical factor for the protection conferred and no preferential effect was observed when sympatric multi-strains mixes were used. This work demonstrates the potential of leveraging the oyster microbiota to enhance long-term disease resistance in oysters and sheds light on the importance of considering the host origin in such protective mechanisms.

Targeting early developmental stages as a strategic window for probiotic application to induce long-term protection has been proposed and explored in various animal models such as mammals or humans (see review by Hashemi *et al*. 2016), but also those relevant to livestock production (Wang et al. 2022; Villumsen et al. 2023). Introducing beneficial microorganisms during these stages can influence both the host’s microbiota composition and immune system development, potentially leading to long-term beneficial immunomodulation. Our results are in line with these findings since we observed a shift in both the transcriptional pattern and microbiota composition of oysters exposed to beneficial microorganisms compared to their non-exposed counterparts, even seven months after the exposure. The long-lasting transcriptional responsiveness was found to be influenced by the host’s origin and was specific to the type of microbial treatment administered. A significant portion of the differentially expressed genes in exposed oysters were associated with immune functions, with a particular emphasis on pattern recognition receptors (PRRs). Intriguingly, the observed difference in phenotype between oysters stimulated with the whole microbiota and those stimulated with multi-strain bacterial mixes could not be fully explained through a thorough analysis of the differentially expressed genes. This suggests that additional factors or intricate interactions within the oyster’s immune system and microbiota may contribute to the differential response. Furthermore, when the oysters were challenged with OsHV-1 µVar three hours after exposure, changes in the transcriptional pattern were still evident in oysters exposed to beneficial microorganisms compared to their non-exposed counterparts, albeit to a lesser extent than before the infectious challenge. This indicates a dynamic interplay between the immune response against the virus and the prior microbial stimulation, with the virus potentially exerting a more pronounced effect on the transcriptional response.

Our findings further indicates that exposure to either bacterial mixes or whole microbiota, leads to changes in the microbiota composition. This was observed during the exposure but also on a long-term basis as previously observed in other studies (Padeniya et al. 2022; Villumsen et al. 2023; Takyi et al. 2024). Interestingly, the bacteria added as part of the bacterial mixes were not detected using the employed method. This suggests that the added bacteria did not effectively integrate the oyster microbiota, even shortly after the start of the exposure. Similar studies indicate that administered bacteria fail to establish and only persist temporarily in the microbiota of exposed animals. For instance, the *Aeromonas sp.* strain administered to oyster larvae was undetectable 72 hours after addition (Gibson et al. 1998). Similarly, exposing the European abalone (*Haliotis tuberculata*) to the *Pseudoalteromonas sp.* hCg-6 exogenous strain resulted in a transient establishment of the probiotic strain in the haemolymph rather than a sustained interaction (Offret et al. 2018). Additionally, Arctic Char (*Salvelinus alpinus*) exposed to various probiotic strains did not show detectable levels of the administered strains four weeks after probiotics administration (Knobloch et al. 2022). The change in microbiota composition observed on long term basis might thus be linked to ongoing interactions between the microbiota and the immune system, leading to a continuous reshaping of both elements and explaining also the observed long-term transcriptional changes.

## Conclusion

Our study successfully investigated methods which aimed at exposing oysters to specific beneficial microorganisms during larval rearing to educate their immune system. We took into account the potential risks associated to this microbial exposure while ensuring that the oysters’ innate immune system was primed for improved disease resistance. We demonstrated the potential of leveraging this microbial education to enhance disease resistance to two major oyster pathogens, OsHV-1 µVar and *V. aestuarianus*, which are current critical threat for oyster farming worldwide. Additionally, our findings emphasize the potential of using controlled whole microbiota transfers as the best strategy to safeguard oyster health in aquaculture settings. Additional optimizations will be required to identify the most effective settings for enhancing the beneficial impact of microbial education. The timing, duration of exposure, and rearing conditions are essential factors for the practical application of this approach in aquaculture environments. Exploring combinations with other strategies, such as selecting oysters with genetic backgrounds that are more receptive to microbial education, is another avenue that certainly deserves further investigation.

## List of abbreviations

ASV: Amplicon Sequence Variants
CFU: Colony–forming unit
DEG: Differentially expressed gene
HR: Hazard-Ratio
LEfSe: Linear discriminant analysis (LDA) Effect Size
MB: Marine Broth
NSI: Naissains Standardisés Ifremer or standardised Ifremer spats
OsHV-1 µVar: Ostreid Herpes Virus 1 µVar
OTU: Operational Taxonomic Unit
pf: post-fertilization
POMS: Pacific Oyster Mortality Syndrome
RNA–Seq: Sequencing of the polyadenylated ribonucleic acids

## Acknowledgments

The authors warmly thank the staff of the Ifremer stations of Argenton, La Tremblade and Arcachon for their help and hospitality during the various oyster sampling campaigns. We are grateful to Leo Duperret, Emily Kunselman, Nicole Faury, Cyrielle Lecadet and Delphine Tourbiez for their help during the oyster experimental infections and Abdellah Benabdelmouna and Christophe Ledu for their help during the larval rearing. We are grateful to Jean-François Allienne, Margot Doberva and Michèle Laudié from the Bio-Environment platform (UPVD, Région Occitanie, CPER 2007-2013 Technoviv, CPER 2015-2020 Technoviv2) for technical support in library preparation and sequencing. We are grateful to the BIO2MAR platform (http://bio2mar.obs-banyuls.fr) for access to instrumentation.

## Authors’ contributions

LDa, LDé, BM, BP, EM, GC and JVD contributed to oysters sampling. LDa, PC, RL and LI contributed to bacteria collection. LDa, PC, LDé, BM, BP, MM, EM, JVD, ET and CC performed oyster experiments. LDa, PC, JFA, CG, OR, JVD, ET and CC prepared samples and performed DNA and RNA extraction on oysters samples for analyses. LDa, MAT, JFA, OR and JP performed qPCR analyses. LDa, ET and CC performed microbiota analyses. LDa, JVD, ET and CC performed RNAseq analyses. LDa, LDé, BM, MAT, BP, MM, YG, JVD, ET and CC conceptualized and designed the experiments. LDa, LDé, BM, MAT, BP, YG, JVD, ET and CC wrote the original draft. LDa, YG, JVD, ET and CC involved in funds acquisition. All authors read and approved the final manuscript.

## Availability of data and materials

Raw sequence data for RNA-seq and 16S sequencing for metabarcoding analysis have been made available through the SRA database (BioProject accession number PRJNA1078733).

R script for survival, DEseq2 and microbiota analyses are available by using the following link: https://zenodo.org/records/11200726.

## Ethical approval

The animal (oyster *Crassostrea gigas*) testing followed all european regulations concerning animal experimentation. The authors declare that the use of genetic resources fulfill the French and EU regulations on the Nagoya Protocol on Access and Benefit-Sharing (French legislation 2019-486).

