## Supplementary_File_2 for "Microbial education plays a crucial role in harnessing the beneficial properties of microbiota for infectious disease protection in *Crassostrea gigas*"

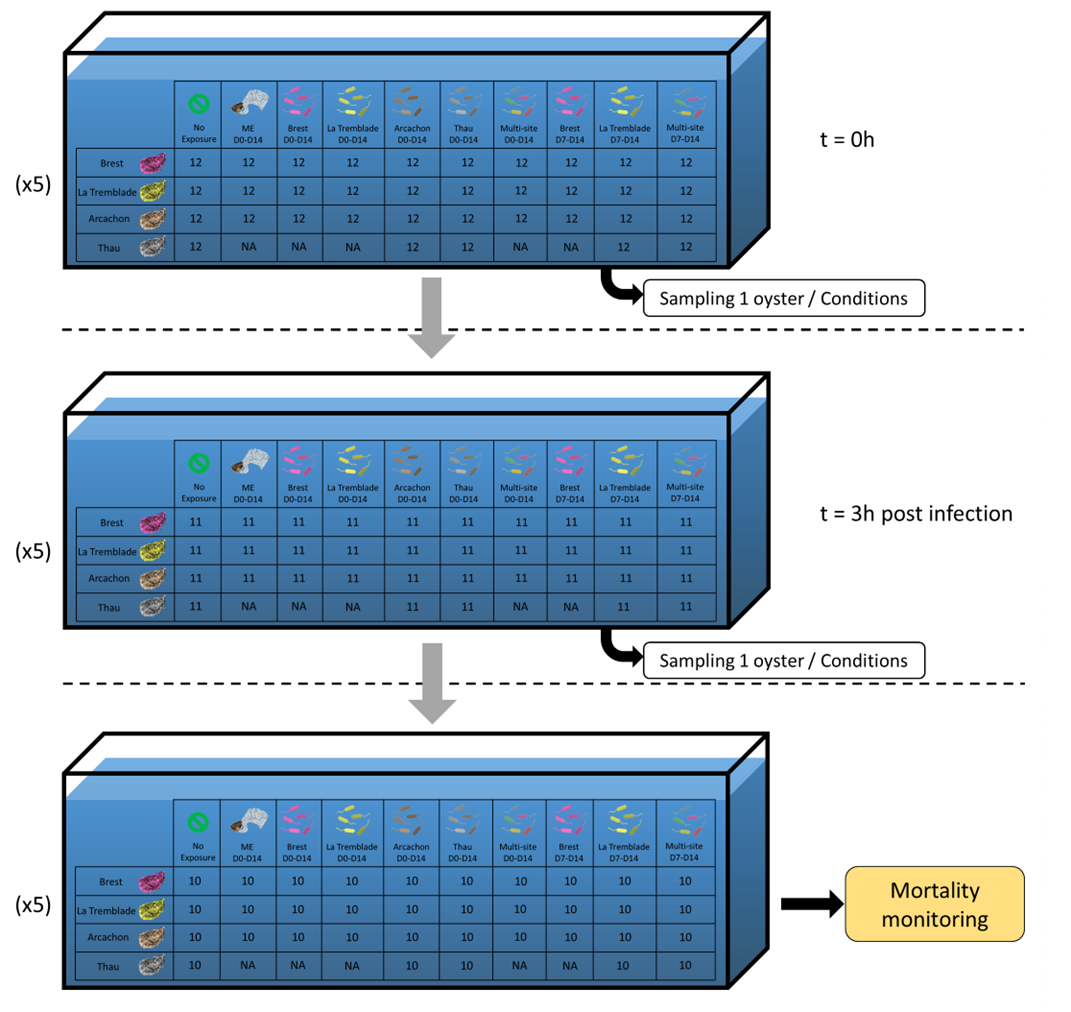

**Figure S1: Experimental design for OsHV-1 experimental infection.**

Prior to OsHV-1 µVar infection (t = 0h), twelve oysters of each population exposed to each microorganism exposure conditions were put in a 50L tank filled with filtered and UV-treated seawater and maintained at 20°C with adequate aeration. One oyster of each population exposed to each microorganism exposure conditions was sampled Before the infection (t = 0h) and three hours after the beginning of the OsHV-1 µVar infection (t = 3h post infection). The oysters sampled were grounded in liquid nitrogen to a powder stored at -80°C and then used for DNA and RNA extraction. After the last sampling, the experimental infection continued, and a mortality monitoring was performed during eight days.

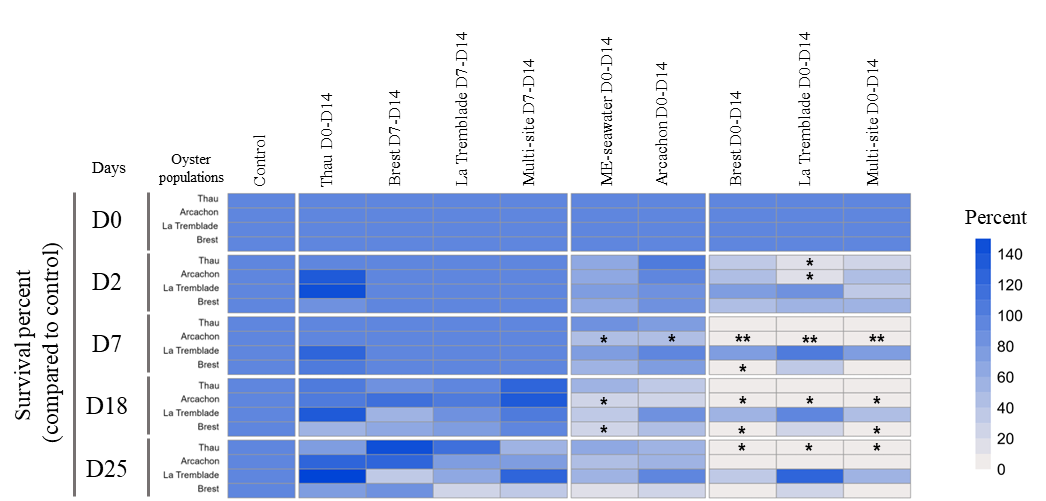

**Figure S2: Multi-strain bacterial mixes displayed contrasted effects on larval survival.**

The effects of the exposure on the survival of larvae from the four oyster populations at different times of the larval rearing stages were recorded. The heatmap shows the survival (in percent) of the larvae exposed to microorganisms compared to the control condition. The asterisks (*) inside the box indicate statistical differences compared to control condition (* p < 0.05 ; ** p < 0.01 ; *** p < 0.001).

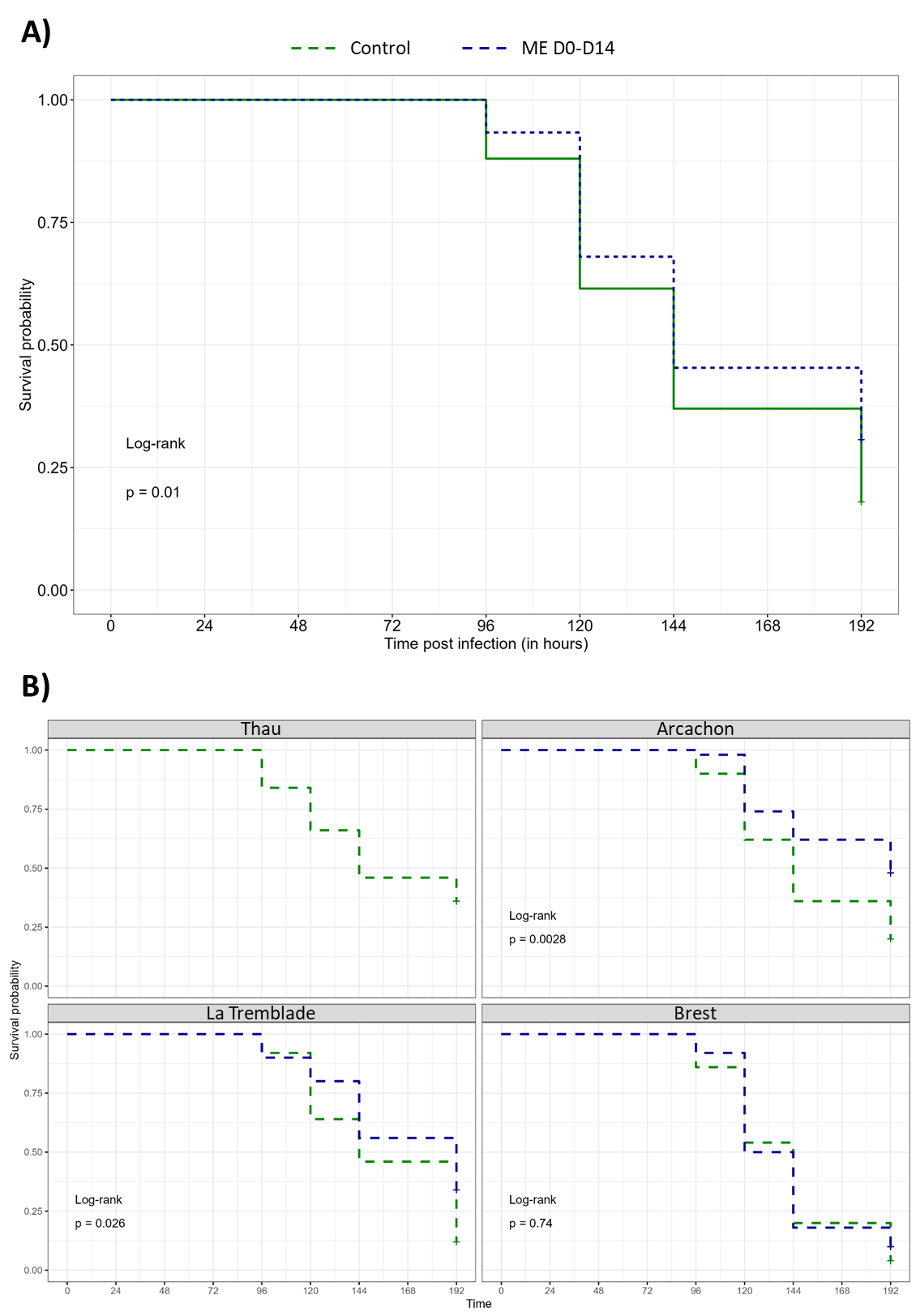

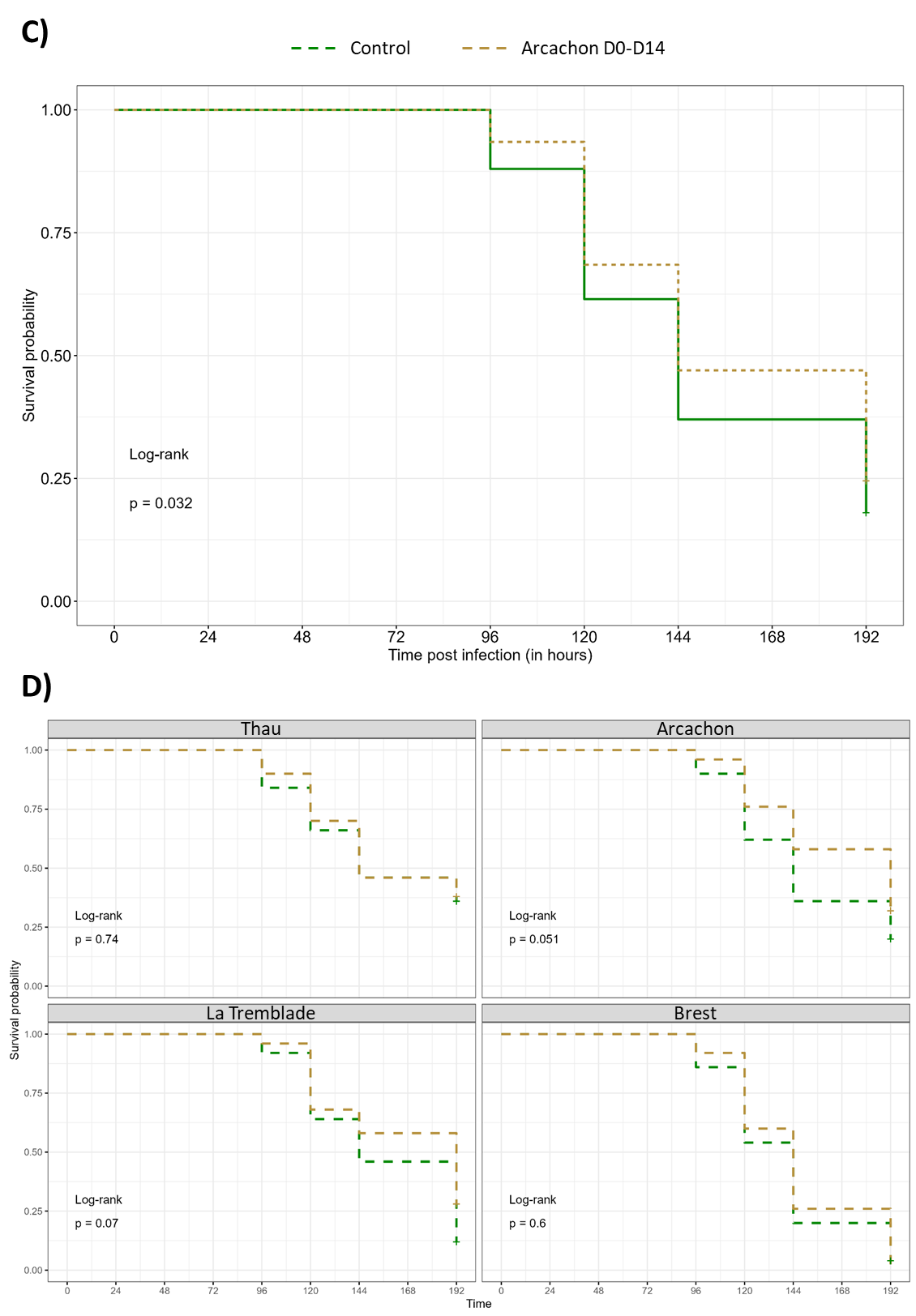

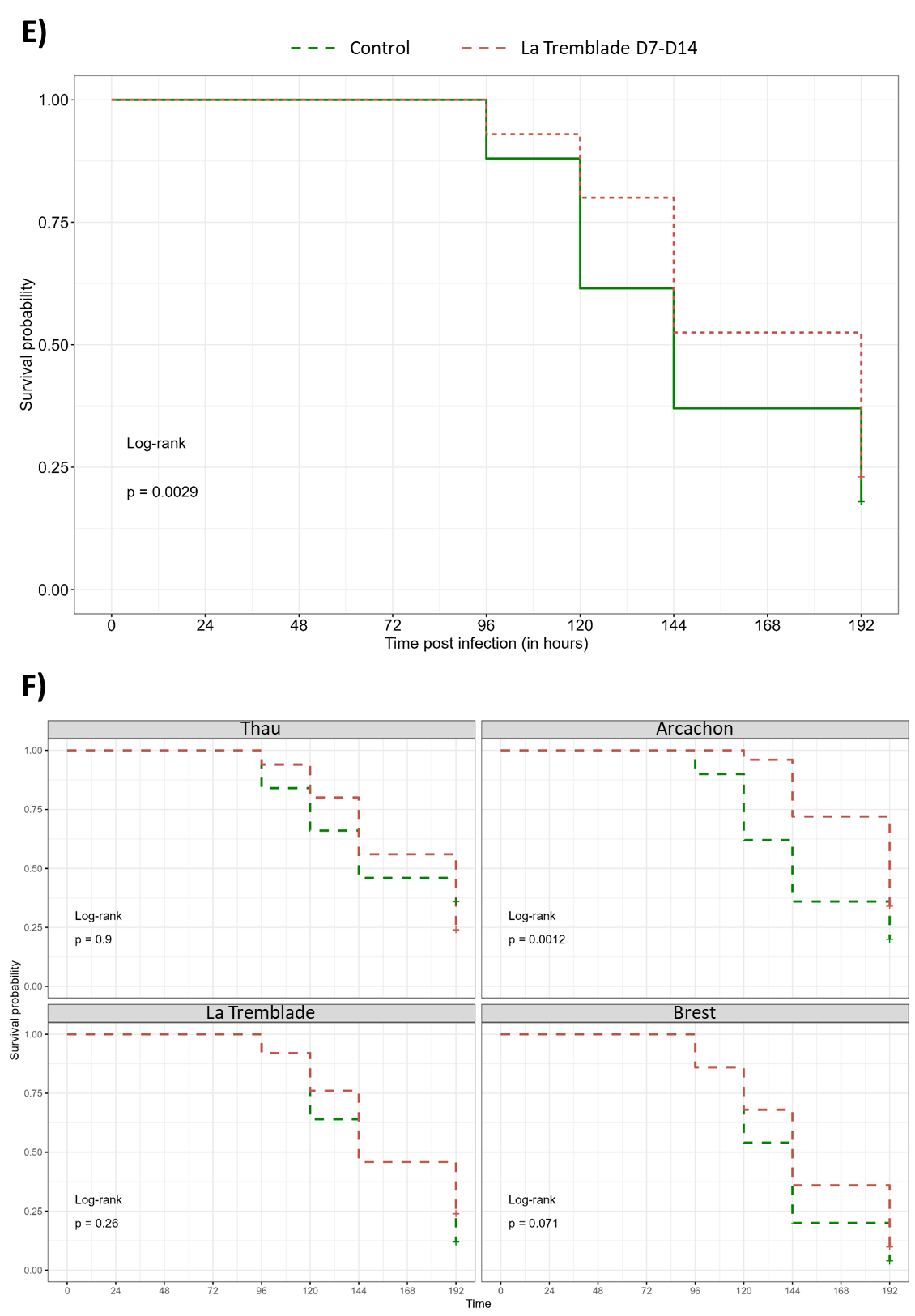

**Figure S3:** **Kaplan-Meier survival curve of OsHV-1 experimental infection.**

Survival curves for oysters exposed: to the microbiota of donor oysters (ME) for all populations combined (n= 150) (A) and details of each population (n= 50 per population) (B); to the Arcachon mix between D0 and D14 (Arcachon D0-D14) for all populations combined (n = 200) (C) and details of each population (n= 50 per population) (D); to the La Tremblade mix between D7 and D14 (La Tremblade D7-D14) for all populations combined (n = 200) (E) and details of each population (n= 50 per population) (F).

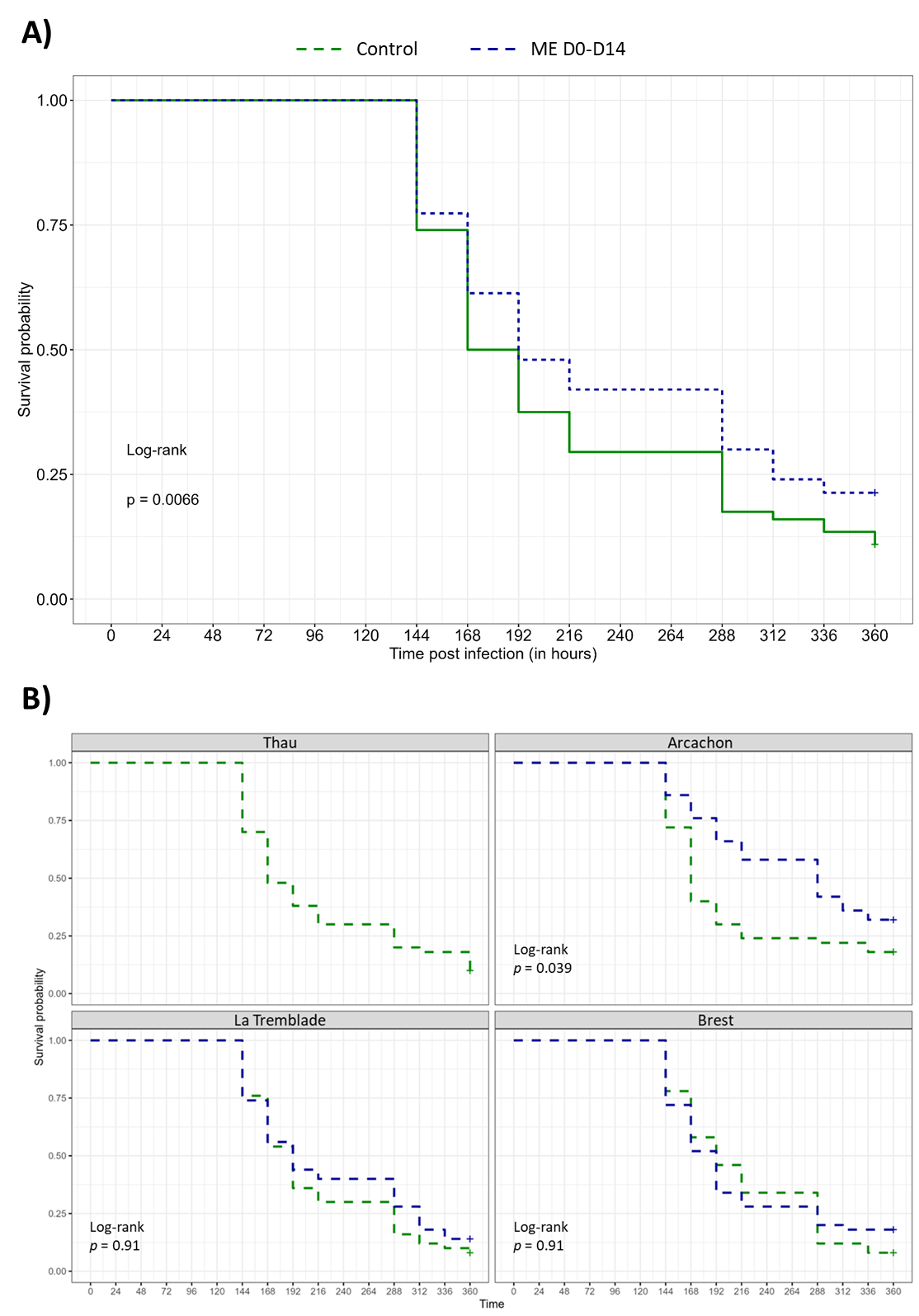

**Figure S4:** **Kaplan-Meier survival curve of *Vibrio aestuarianus* experimental infection.**

Survival curves for oysters exposed: to the microbiota of donor oysters (ME) for all populations combined (n=150) (A) and details of each population (n= 50 per population) (B).

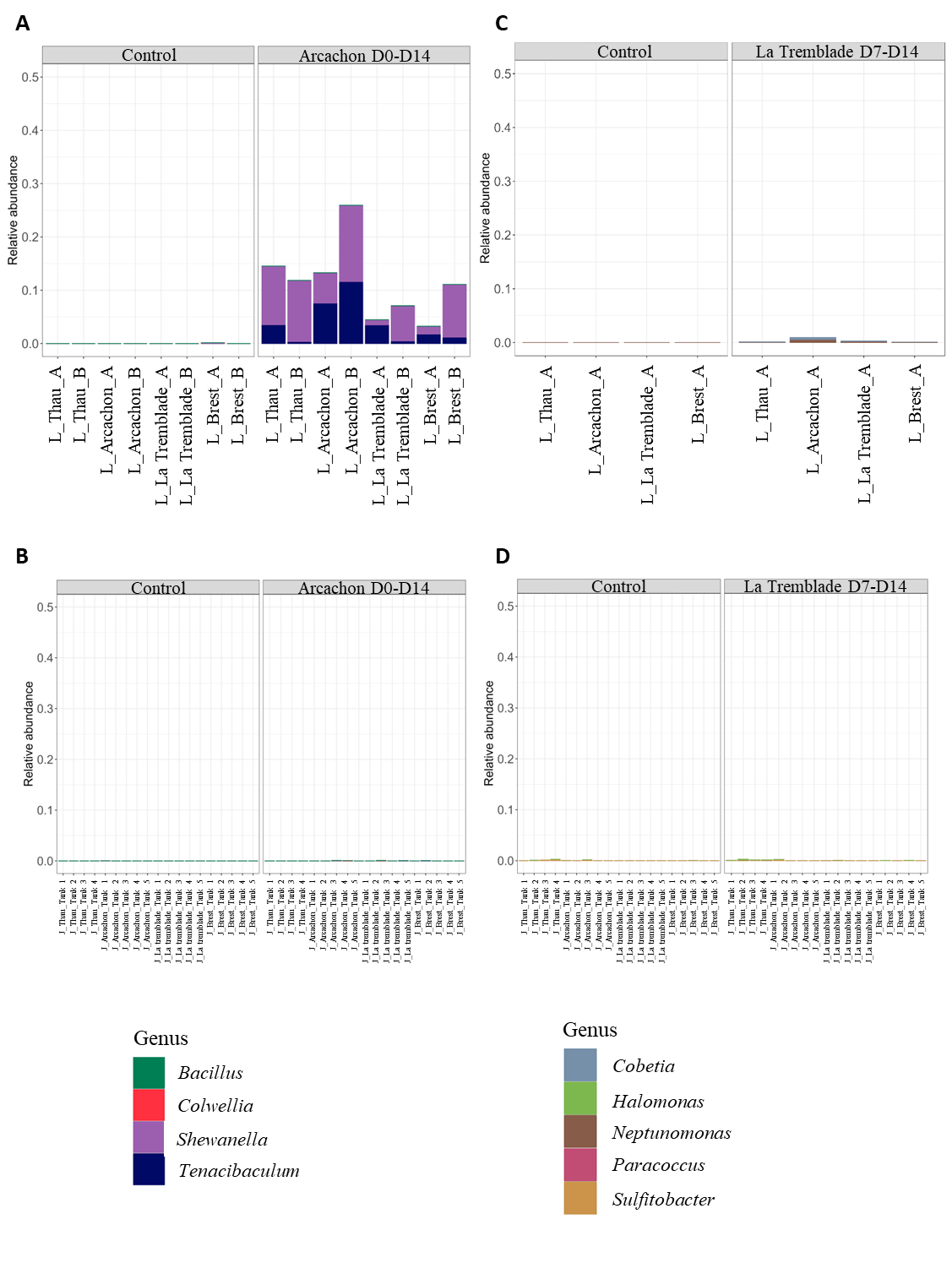

**Figure S5: Distribution at the genus level of bacteria administered by the Arcachon mix and La Tremblade mix.**

Distribution of bacteria administered by the Arcachon mix during the larval rearing step after seven days of exposure (A) or during the juvenile stages seven month after the exposure (B) and distribution of bacteria administered by the La Tremblade mix during the larval rearing step after seven days of exposure (C) or during the juvenile stages seven month after the exposure (D). For larval stages (A) and (C), analyses were performed on each oyster population (indicated on x-axis) on 2 pools of 10000 larvae sampled in 2 independent tanks for exposure to Arcachon D0-D14 (A) and on 1 pool of 10000 larvae sampled in a unique tank for exposure to La Tremblade D7-D14 (B). For juvenile stages (B) and (D), analyses were performed on 4 to 5 individuals of each population sampled in independent tanks (indicated in x-axis).

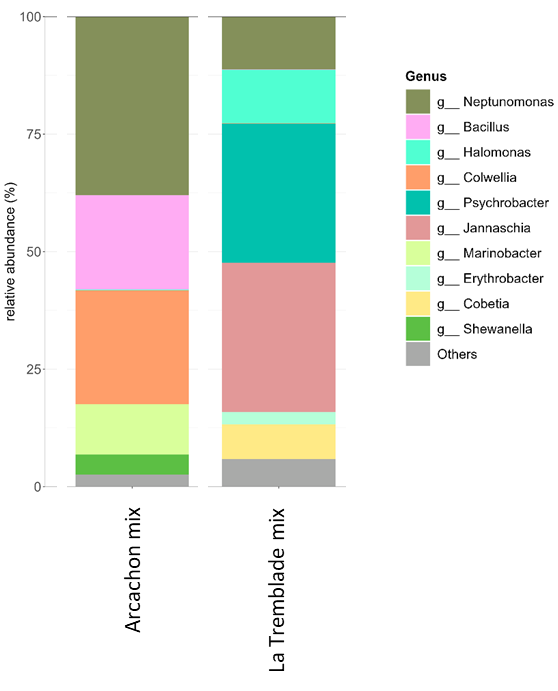

**Figure S6: Distribution at the genus level of bacteria composing the mock control for Arcachon and La Tremblade mixes.**
